# *Candida albicans* promotes self-adhesion and tumor cell progression of colorectal cancer cells

**DOI:** 10.64898/2026.09.09.750446

**Authors:** Janet F. Staab, Olivia K. A. Paulin, Julian R. Naglik, Nicholas C. Zachos

## Abstract

*Candida albicans*, a benign commensal yeast in the healthy gut, is increasingly being recognized as a tumor-promoting component of the gut mycobiome that is linked to worse patient outcomes in colorectal cancer (CRC). Emerging evidence suggests that in the tumor microenvironment, *C. albicans* drives tumor progression in its pathogenic hyphal form with the secretion of its peptide toxin, candidalysin, enhancing tumor cell evolution. CRC cells differentially express transglutaminase 2 (TG2), which mediates covalent binding to *C. albicans* and can promote tumor adaptability and metastases. However, mechanistic insight into how *C. albicans* infection leads to CRC progression is incomplete. CRC lines that express high (SW480) and low (LoVo, HCT 116) TG2 were infected with *C. albicans* and examined for effects on epithelial to mesenchymal transition (EMT) and cell migration. Parallel studies were also conducted using a candidalysin-deficient *C. albicans* strain. *C. albicans* invasion stimulated TG2 activity that allowed covalent crosslinking of the hyphae adhesin, Hwp1, to SW480 cells in a candidalysin-dependent manner. Independent of TG2 expression, infection of SW480, LoVo, and HCT 116 cells induced EMT-like changes including downregulation of E-cadherin, altered *N*-glycosylation of N-cadherin, and variable Snail1 expression. Infection increased MGAT5 activity, which raised levels of branched β(1-6) *N*-glycans. Infection promoted CRC cell motility that was blocked by pharmacological inhibition of MGAT5. These findings reveal that *C. albicans* can promote covalent attachment to TG2-high CRC cells and induce EMT-like and migration-associated changes through TG2-independent alterations in epithelial plasticity and *N*-glycosylation.

## Introduction

Colorectal cancer (CRC) is a leading cause of cancer mortality worldwide. Environmental contributors such as diet, obesity, and smoking are implicated in CRC cases that lack a strong inherited basis (1, 2). These factors reshape the gut microbiome with growing evidence linking microbiome alterations to CRC pathogenesis (3–6). Although bacteria have been the primary focus, recent metagenomic studies also implicate the mycobiome: *Candida* species, especially *C. albicans*, are enriched in gastrointestinal tumors and associate with worse outcomes (7, 8).

*C. albicans* commonly resides in the oral and gut mucosa as a commensal yeast but can switch to a filamentous (hyphal) form that adheres to and invades tissues. Hyphae express adhesins (9–12) and secrete candidalysin, a peptide toxin that damages the epithelium and triggers inflammation (13–16). In oral squamous cell carcinoma, invasive *C. albicans* induces an epithelial-to-mesenchymal transition (EMT) program, characterized by E-cadherin loss, N-cadherin upregulation, and increased migratory behavior (17), supporting fungal contributions to tumor progression. Whether *C. albican*s assumes invasive morphologies in colorectal tumors and how fungal/tumor interactions affect CRC cell behavior have not been defined.

Transglutaminase 2 (TG2) is a multifunctional enzyme implicated in cell survival, extracellular matrix remodeling, and EMT-related signaling (18–22). TG2 expression and activity correlate with advanced CRC (23–26). Transglutaminases covalently crosslink microbial substrates to host proteins (12, 27, 28) including the fungal Hyphal wall protein 1 (Hwp1), a known transglutaminase substrate that mediates stable fungal attachment to oral epithelia (12, 29, 30). Elevated TG2 in CRC tissues could therefore create an enzymatic niche facilitating covalent attachment and persistence of *C. albicans* hyphae at tumor sites.

From this knowledge, we set out to determine the impact of *C. albicans* infection in human CRC progression and whether TG2 activity mediates phenotypic changes in CRC cells. We tested the hypothesis that *C. albicans* induces TG2 activity and promotes EMT in CRC cells. To do this, we evaluated the mechanisms governing (1) adherence of *C. albicans* with CRC cells, (2) activation of EMT programming, and (3) increased migration in *C. albicans*-infected CRC lines (i.e., SW480, HCT 116, LoVo). From our studies, we found that invasive *C. albicans* activated TG2 in a candidalysin-dependent manner and promoted crosslinking of recombinant Hwp1 in TG2-high expressing CRC cells (i.e., SW480), supporting a mechanism for stable fungal adherence. *C. albicans* also invaded TG2-low expressing CRC cells (i.e., HCT116, LoVo) indicating Hwp1-independent fungal adhesion occurred. Concurrently, invasive *C. albicans* induced an EMT-like program independent of TG2 expression levels characterized by the loss of E-cadherin with increased branched *N*-glycosylation of N-cadherin. Cell migration was enhanced in all CRC cells exposed to *C. albicans* that was suppressed by blocking complex *N*-glycosylation in SW480 cells. Importantly, *Candida*-induced EMT features persisted when TG2 activity was inhibited, further supporting that fungal modulation of tumor cell plasticity occurred independently of TG2-mediated adhesion.

Together, these results support a dual mechanism model in which *C. albicans* uses complementary strategies to influence CRC cells: (1) candidalysin-and TG2-dependent covalent attachment that promotes fungal colonization at tumor sites; and (2) candidalysin-and TG2-independent signaling and post-translational modifications of host proteins that drive EMT-like changes and increased cell migration. Framing the study around these two mechanisms clarifies how fungal colonization directly modulates CRC tumor cell evolution through epithelial plasticity and N-glycosylation-mediated migration.

## Materials and Methods

### Candida strains

Reference strain, SC5314 (31), was used in all experiments unless otherwise indicated. Some experiments used the SC5314 derivative (AHY940) deleted for the *ECE1* gene encoding candidalysin (*ece1*∆/∆; OP02) or the reconstituted strain (*ece1*∆/∆ +*ECE1*; OP17) (32). Briefly, candidalysin mutant strains were generated using a CRISPR-Cas9 gene deletion method described previously (33). AHY940 (SC5314 *leu2*Δ/*LEU2*) is a direct descendent of SC5314, heterozygous at the *LEU2* locus, that facilitates CRISPR cassette integration and recycling. The *ece1*Δ/Δ strain (*leu2*Δ/*LEU2 ece1*Δ/Δ) is an *ECE1* open reading frame deletion mutant constructed in parental strain AHY940. The *ece1*Δ/Δ+*ECE1* re-integrant strain (*leu2*Δ*/LEU2 ece1*Δ/Δ+*ECE1)* was constructed in the *ece1*Δ/Δ strain. Other *Candida* species, *C. tropicalis*, *C. krusei*, *C. auris,* and *C. glabrata* were obtained from the Clinical Microbiology Laboratory at the Johns Hopkins Hospital. The organisms were stored at -80°C and cultured in liquid or agar plates of yeast extract (2%) peptone (2%) dextrose (1%) (YPD) medium at 30°C according to standard methods (34).

### Tissue culture lines

The colorectal cancer lines SW480 and HCT 116 were obtained from American Type Culture Collection (ATCC; #CCL-228 and CCL-247, respectively). LoVo (ATCC #CCL-229) cells were a kind gift from Bhuminder Singh at Vanderbilt University Medical Center. SW480 cells were maintained in Leibovitz’s L-15 Medium (Thermo Fisher Scientific, #11415064) with 10% fetal bovine serum (FBS) and 1% penicillin/streptomycin (Thermo Fisher Scientific, #15140122) (complete L-15 medium) at 37°C in the absence of CO2 per ATCC recommendations. HCT 116 were maintained in McCoy’s 5A (Modified) Medium (Thermo Fisher Scientific, # 16600082), 10% FBS and 1% penicillin/streptomycin. LoVo cells were maintained in Ham’s F-12K (Kaighn’s) Medium (Thermo Fisher Scientific, # 21127022), 1% Glutamax (Thermo Fisher Scientific #35050061), 10% FBS and 1% penicillin/streptomycin at 37°C in a humidified CO_2_ incubator. For experiments, subconfluent cultures (80-90%) were harvested by trypsin digestion and the lifted cells counted using an automated counter (Countess II; Invitrogen). Cells (2-5 X 10^5^) were seeded in 12-well plates with sterile cover slips or in 4-chamber slides (Lab-Tek, #154526) (for *in situ* TG2 activity and immunofluorescence) or in 12-well plates (for western blot analysis) and allowed to reach confluence (∼5 X 10^5^/well) for 3-5 days prior to infection with *Candida* yeasts.

### Human colonic organoids

Patient-derived organoids (PDOs) were derived from healthy donors undergoing routine colonoscopies at the Johns Hopkins Hospital using our established protocol (35, 36). De-identified patient tissues were obtained with consent under the Johns Hopkins University IRB protocol #NA_00038329, and all methods were performed according to approved guidelines and regulations. PDOs were propagated as 3D spheroids in 25 μL Matrigel (Corning, #356231) domes in Wnt3a-, Rspo1-, and Noggin-containing media (37). To establish monolayers, 3D PDOs were fragmented using TrypLE Express (Thermo Fisher Scientific, #12604013) (36) and seeded onto collagen IV-coated 0.4 μm polyester membrane Transwells (Corning, #3470). PDO monolayers were grown to confluency and then exposed to media lacking Wnt3a and R-spon1 for 5 days (35, 36) to induce differentiation of mature epithelial cell types (e.g., goblet cells). At day 5 of differentiation, the monolayers generated from two donors (n=2) were used for *C. albicans* infections followed by 5-BPA crosslinking reactions.

### Infection of CRC cells

*Candida* yeasts were grown in YPD at 30°C with shaking for 22-24 h. Yeasts were washed with sterile phosphate-buffered saline, pH 7.4 (PBS) and adjusted to 5 X 10^6^/mL in complete appropriate media. Wells with near confluent cultures were washed twice with Hank’s Balanced Salt solution (HBSS) with calcium and magnesium (Corning, #21-023-CV) and 2.5 X 10^5^ yeasts added in a 400 μL final volume to achieve a multiplicity of infection (MOI) of 2 (i.e., two yeasts/CRC cell) for 3 h at 37°C. Some *C. albicans* infections were extended to 6 or 16 h (MOI of 0.5 for 16 h infections) to examine the effect on epithelial to mesenchymal transition marker proteins by western blot analysis.

### Infection of patient-derived organoid monolayers

*C. albicans* yeasts grown in YPD at 30°C with shaking for 22-24 h were washed with sterile PBS and adjusted to 5 X 10^6^/mL in differentiation medium. The apical media of colonic PDO monolayers was removed and replaced with 100 μL (5 X 10^5^ yeasts; MOI of 5) of *C. albicans* suspension on day 5 of PDO differentiation (35, 36). The co-culture was incubated for 3 h in a 37°C/CO_2_ tissue culture incubator. At the 3 h time point, the co-cultures were used for in-situ transglutaminase activity assays.

### *In situ* transglutaminase 2 (TG2) activity assay

Activation of TG2 enzymatic activity on the surface of SW480, HCT 116, or LoVo cells was visualized by cross-linking 5-(biotinamido)pentylamine (5-BPA) (Sigma-Aldrich, #914134) or recombinant Hwp1HA (see generation of rHwp1HA below). Post infection or wounding by scratch assay, CRC cells were washed twice in HBSS without calcium or magnesium (Corning, #21-021-CM) and incubated for 1 h in 400 µL reaction buffer (100 mM Tris-HCl pH 7.5, 10 mM CaCl_2_, 1 mM DTT, 1 mM EDTA, 5 µM 5-BPA or 2.5 µM rHwp1HA) at 37°C. To validateTG2 activity, parallel reactions were incubated with Z-DON (Sigma-Aldrich, #616467), a cell-permeant, TG2-specific activity inhibitor. In these experiments, SW480 monolayers were preincubated with 100 µM Z-DON for 1 h prior to and during the TG2 activity assay. To determine whether TG2 activation required an influx of calcium (Ca^2+^), SW480 monolayers were preincubated with 10 μM BAPTA-AM (Sigma-Aldrich, #196419) for 30 min at 37°C prior to 3 h *C. albicans* infection. BAPTA-AM was also included in the post treatment TG2 activity assay. Crosslinking reactions were stopped by washing the cells twice with PBS and fixation with 4% paraformaldehyde in PBS on ice for 15 min Fixed cells were stored at 4°C in PBS until used for 5-BPA or rHwp1HA detection and immunofluorescence.

### *In situ* transglutaminase 2 (TG2) activity assay in human patient-derived organoids

At the end of the 3 h *C. albicans* infection period, the apical and basolateral surfaces of PDO monolayers were washed twice with warm HBSS (no calcium or magnesium) followed by incubation in TG2 reaction solution with 50 μM 5-BPA for 15 min at 37°C/CO_2_. The reaction was stopped by washing the monolayers 2X with PBS and fixation in 4% paraformaldehyde in PBS at room temperature for 30 min Fixed monolayers were stored 4°C in PBS until 5-BPA detection and immunofluorescence.

### Detection of in-situ incorporation of 5-BPA and immunofluorescence

Fixed CRC cells infected with *Candida* or treated with candidalysin peptide (see below) followed by 5-BPA cross-linking reactions were blocked in 15% fetal bovine serum, 2% bovine serum albumin in PBS (blocking buffer) for 30 min After washing with PBS, the cells were incubated with rabbit antiserum to *C. albicans* (Abcam, #AB53891) and streptavidin-AF647 (Molecular Probes/Thermo Fisher Scientific, #S32357) both at 1:200 in PBS for 30 min at room temperature. The antiserum also bound to other *Candida* species tested for activation of TG2 in SW480 cells. Incubation with *C. albicans* antiserum was omitted in monolayers wounded by scratching or candidalysin exposure, and in monolayers incubated with rHwp1HA in the crosslinking reaction. After washing with PBS, treated CRC cells were incubated with goat anti rabbit IgG conjugated to AF488 (Thermo Fisher Scientific, # A11008) at 1:200 in PBS for 30 min In some instances, washed cells were permeabilized with 0.5% Triton X100 in dH_2_O for 30 min at room temperature followed by incubation with phalloidin conjugated to AF568 (1:100) (Molecular Probes/Thermo Fisher Scientific #A12380) in PBS for 30 min at room temperature. The cover slips were subsequently washed twice with PBS and the nuclei stained with Hoechst 33258 (Sigma-Aldrich, #94403) at 1:5000 in PBS for 15 min. Washed coverslips or chambers were mounted in ProLong Gold (Invitrogen/Thermo Fisher Scientific #P36930) and allowed to cure at room temperature for 18-24 h. Incorporation of 5-BPA was observed under confocal microscopy (Olympus FV3000RS) and 5-BPA+ cells enumerated in 9 ROIs at 20X magnification. This staining scheme detects extracellular fungal surfaces; internalized fungal elements (invading hyphae) appear dark.

### Detection of *in situ* incorporation of 5-BPA and immunofluorescence of patient-derived organoids

Fixed PDOs were simultaneously permeabilized and blocked in 15% fetal bovine serum, 2% bovine serum albumin, 0.1% saponin (Sigma-Aldrich, #S4521) in PBS for 30 min at room temperature. The monolayers were washed with PBS and incubated with rabbit antiserum to *C. albicans* at 1:200 in block solution for 30 min at room temperature. After washing 2X with PBS, PDOs were incubated with goat anti-rabbit IgG conjugated to AF488 (1:100), streptavidin conjugated to AF647 (1:200) and phalloidin conjugated to AF568 (1:100) in PBS for 30 min at room temperature. The monolayers were washed 3X with PBS and incubated with Hoechst 33258 at 1:5000 in PBS for 15 min followed by one PBS wash. The Transwell membranes were cut away from the insert and mounted in ProLong Gold and allowed to cure at room temperature for 18-24 h before imaging by confocal microscopy (Olympus FV3000RS).

### Detection of *in situ* incorporation of rHwp1HA and immunofluorescence of SW480 cells

Recombinant Hwp1HA was used as a surrogate for *C. albicans* hyphae in TG-mediated attachment to SW480 cells. SW480 monolayers grown in 4-chambered slides were infected with *C. albicans* for 3 h (MOI 2) followed by *in situ* TG2 activity assays. The TG2 reaction buffer contained 2.5 µM rHwp1HA in place of 5-BPA and the chambered slides were incubated for 1 h at 37°C. Parallel reactions had 100 µM Z-DON to inhibit TG2 activity or 50 μg/mL dextran sulfate to compete with candidalysin (38). Washed monolayers were fixed in 4% PFA on ice for 15 min and stored in PBS at 4°C until detection of crosslinked rHwp1HA. After washing twice with PBS, the cells were blocked in for 30 min and incubated with rabbit *C. albicans* antiserum and mouse anti-HA conjugated to AF488 (Invitrogen, #A21287), both at 1:200 dilution, in blocking buffer for 30 min at room temperature. Following incubation with primary antibodies and washing, in some experiments, the cells were permeabilized with 0.5% Triton X-100 at room temperature for 30 min. The chamber slides were washed and incubated with goat anti-rabbit IgG conjugated to AF568 (Invitrogen/Thermo Fisher Scientific A-11011) at 1:100 dilution and phalloidin-AF633 (1:200) (Invitrogen/Thermo Fisher Scientific, # A22284) in PBS for 30 min After incubation with fluor-labeled reagents, the cells were washed twice with PBS and the nuclei stained with Hoechst 33258 for 15 min. After a final wash with PBS, the chambers removed from the slide and the cell monolayers mounted in ProLong Gold (Molecular Probes/Thermo Fisher Scientific #P10144). The samples were allowed to cure at room temperature in the dark for 18-24 h before analysis by confocal microscopy. Incorporation of rHwp1HA was observed under confocal microscopy (Olympus FV3000RS) and rHwp1HA+ cells enumerated in 9 ROIs at 20X magnification. This staining scheme detected extracellular fungal surfaces; internalized fungal elements appear dark.

For sequential crosslinking of 5-BPA and rHwp1HA, SW480 monolayers grown in 4-chambered slides were infected with *C. albicans* for 3 h (MOI 2) followed by two in-situ TG2 activity assays. Control cells did not receive *C. albicans* yeasts. After removing unbound hyphal cells by washing twice in HBSS without calcium or magnesium, the cells were incubated in 400 μL TG2 reaction buffer with 5 μM 5-BPA for 30 min at 37°C. The cell monolayers were washed with HBSS (no calcium or magnesium) and incubated in 400 μL TG2 reaction buffer with 2.5 μM rHwp1HA for 30 min at 37°C. At the end of the second 30 min incubation, the cells were washed twice with PBS and fixed on ice in 4% PFA for 15 min. *C. albicans* was detected with concanavalin A conjugated to TRITC (Invitrogen, #C860; (39, 40)) at 5 μg/mL in PBS for 15 min protected from light. Concanavalin A binds to high mannose residues and will preferentially react with highly mannosylated fungal cell walls. After washing with PBS, the cells were blocked in 15% fetal bovine serum, 2% bovine serum albumin in PBS for 30 min. After washing with PBS, the cells were treated with mouse anti HA antibody conjugated to AF488 (1:100) in blocking buffer for 30 min. The slide chambers were washed twice with PBS and the cells incubated with streptavidin-AF647 (1:200) in PBS for 30 min followed by three PBS washes and incubation with Hoechst (1:5000) in PBS for 15 min. The slide chambers were removed and the cells mounted in ProLong Gold for confocal analysis. 5-BPA+ and rHwp1HA+ cells were imaged (Leica SP8) and enumerated in 12 ROIs at 20X magnification.

### Immunofluorescence of EpCAM post *C. albicans* infections and 5-BPA or rHwp1HA incorporation on CRC cells

Fixed SW480 monolayers grown in 4-chambered slides infected with *C. albicans* and subjected to TG2 activity assays with 5-BPA were first stained for *C. albicans* extracellular fungal elements with concanavalin A conjugated with TRITC in the dark as above. The cell chambers were washed twice with PBS and incubated in blocking solution for 30 min protected from the light. After washing with PBS, the cells were incubated in blocking solution with antibodies to EpCAM (mouse IgG, Cell Signaling # 2929) for 1 h. After two washes with PBS, the cells were incubated with goat anti mouse IgG conjugated to AF488 (Invitrogen/Thermo Fisher Scientific # A-11001) in block solution at 1:100 plus streptavidin conjugated to AF647 at 1:200 to detect incorporated 5-BPA for 30 min protected from light. The cells were washed twice with PBS, followed by incubation with Hoechst nuclear stain for 15 min. After a final wash, the chamber was removed and the stained cells mounted in ProLong Gold. After curing, the cells were imaged by confocal microscopy (Olympus FV3000RS) and 9 ROIs at 20X magnification captured for *C. albicans* and cell marker staining analyses.

Fixed HCT 116 or LoVo cells grown in 4-chambered slides, infected with *C. albicans* and subjected to TG2 activity assays, were first incubated in blocking solution for 30 min prior to reacting with *C. albicans* rabbit antiserum and mouse IgG to EpCAM in blocking solution as described above for 1 h. After washing with PBS, cells were incubated in blocking solution with goat anti-rabbit conjugated to AF568 at 1:100, goat anti-mouse AF488 at 1:100 and streptavidin conjugated to AF647 at 1:200 for 30 min protected from light. After a final wash, the chamber was removed and the stained cells mounted in ProLong Gold. After curing, the cells were imaged by confocal microscopy (Olympus FV3000RS) at 20X magnification.

Fixed SW480 monolayers grown in 4-chambered slides, infected with *C. albicans* and subjected to TG2 activity assays with rHwp1HA were first stained for *C. albicans* extracellular fungal elements with concanavalin A conjugated with TRITC, as above. The cell chambers were washed twice with PBS and incubated in blocking solution for 30 min protected from the light. After washing with PBS, the cells were incubated in blocking solution with antibodies to EpCAM (mouse IgG) and rabbit anti-HA tag (Rockland, #27745), both at 1:100 in blocksolution for 1 h. After two PBS washes, the cells were incubated with goat anti mouse IgG conjugated to AF633 (Invitrogen/Thermo Fisher Scientific, A-21050) and goat anti-rabbit AF488 in block solution for 30 min protected from light. After a final wash, the chamber was removed and the stained cells mounted in ProLong Gold. After curing, the cells were imaged by confocal microscopy (Olympus FV3000RS) and 9 ROIs at 20X magnification captured for *C. albicans* and cell marker staining analyses.

### SW480 EpCAM abundance quantitation

SW480 cells infected with *C. albicans* followed by 5-BPA or rHwp1HA crosslinking and stained for EpCAM were analyzed in ImageJ2 (Fiji) for surface staining intensities of 5-BPA+ cells for profile gray values. EpCAM staining was stratified as low (EpCAM-low) with gray values ≤30, EpCAM high (EpCAM-high) with gray values 40-60 and EpCAM very high (EpCAM-vhigh) with gray values >100 pixel intensities. Nine ROI with a mean number of 164 cells (+/- 21.2) per ROI were quantified. The same criteria were used to evaluate rHwp1HA-positive cells post *C. albicans* infection.

### TG2 immunofluorescence

SW480 cells grown on coverslips, infected with *C. albicans* for 6 h (MOI 2), followed by 5-BPA crosslinking were fixed in 4% PFA on ice for 30 min. *C. albicans* extracellular surfaces were reacted with rabbit antiserum to *C. albicans* at 1:200 in block solution for 30 min at room temperature. The cells were washed 2X in PBS and incubated with goat anti rabbit IgG AF488 (1:200) in block solution for 30 min. After washing, the cells were permeabilized in 0.5% Triton X-100, washed with PBS and incubated in blocking solution for 30 min. After washing with PBS, the cells were incubated with a mouse antibody to TG2 (ABCAM, #ab2386) at 1:100 in blocking solution for 30 min. The cells were washed 2X with PBS and incubated in goat anti-mouse IgG conjugated to AF568 (1:100) and streptavidin AF647 (1:200) for 30 min in PBS. After three PBS washes, the cells were incubated in PBS with Hoechst nuclear stain for 15 min. After a final wash, the coverslips were mounted in ProLong Gold and analyzed by confocal microscopy at 20X magnification (Olympus FV3000RS).

### Sulfated glycosaminoglycan immunofluorescence

Fixed SW480, HCT 116, and LoVo cells grown in 4-chambered slides were blocked for 30 min, washed with PBS and incubated with anti-heparan sulfate mouse IgM antibody (AMSBIO #3702551) at 1:100 for 1 h at room temperature. The cells were washed 2X in PBS and incubated with goat anti-mouse IgM and IgG AF488 (Thermo Fisher Scientific, #A10680) at 1:100 in block solution for 30 min at room temperature. After three PBS washes, the cells were incubated in PBS with Hoechst nuclear stain for 15 min. After a final wash, the chambers were removed and the slide mounted in ProLong Gold. Cell staining was analyzed by confocal microscopy at 20X magnification (Olympus FV3000RS).

### Candidalysin sensitivity assays

Candidalysin cytotoxicity was measured as previously described (14, 41) with some modifications. SW480 and LoVo cells (2 x 10^5^/well) were seeded in triplicate in 96-well plates in complete media and grown for two days prior to treatment with 15 μM candidalysin for 24 h. Lactate dehydrogenase (LDH) release was measured in the wells using a Sigma-Aldrich LDH Cytotoxicity Assay Kit (#MAK529). Percent cytotoxicity was calculated from total lysis wells per the manufacturer’s specifications.

### Recombinant Hwp1 production and TG substrate reactivity

The coding region for the N-terminal 148 amino acids of mature Hwp1 (aa 40-188) was amplified from SC5314 genomic DNA as previously described (42) using primers H118P 5’-GG<u>CTGCAG</u>GTTCTTATGATTACTATCAAGAACCATGTGATG and H564HA 5’-**TCA**<u>TGCATAGTCCGGGACGTCATAGGGATAGCCCGCATAGTCAGGAACATCGTATGGGTACCC</u>AATATCTGGGATCCAATCAGTTGG engineered with a PstI site (underlined nucleotides), and a double hemagglutinin tag (HA; double underline nucleotides) at the carboxy-terminus of the recombinant protein that preceded a stop codon (bold). The amplicon was digested with PstI and cloned into pPICZαB (Invitrogen/Thermo Fisher Scientific, #V19520) between the PstI and PmlI sites. A sequence-verified clone was introduced into *Pichia pastoris* strain X-33 (Invitrogen/Thermo Fisher Scientific, #C18000) by electroporation for secreted expression of recombinant Hwp1 (rHwp1HA) as previously described (42). Recombinant Hwp1HA was purified from *P. pastoris* culture supernatants by step-wise ammonium sulfate precipitations (43) and dialyzed against 50 mM Tris-HCl pH 8.0, 1 mM EDTA, to remove the ammonium sulfate. The concentration of rHwp1HA was determined using Beer’s law and absorbance at A205 (44, 45). SDS-PAGE of the concentrated rHwp1HA fraction did not reveal other *P. pastoris* proteins by Coomassie blue staining (not shown).

TG substrate activity for rHwp1HA was confirmed by incorporation of 5-BPA using purified recombinant human TG1 (Zedira, #T009) or TG2 (Zedira, #T002). Briefly, 200 μL reactions (100 mM Tris-HCl pH 7.5, 5 mM CaCl_2_, 1 mM DTT, 1 mM EDTA, 50 µM 5-BPA) with increasing concentrations of rHwp1HA and either 1 U/mL of TG1 or TG2 were incubated at 37°C for 15 min and stopped on ice. Negative controls omitted rHwp1HA in the reactions. 5-BPA cross-linked rHwp1HA was subsequently detected by capture in a 96-well plate: 50 μL of 1:20 dilutions in PBS were added to a 96-well plated coated with rabbit anti-HA antibody (Thermo Fisher Scientific, # 600-401-384S) at 10 μg/mL in PBS. After 1 h incubation, the wells were washed with PBS and incubated with Streptavidin-alkaline phosphatase (Southern Biotech, #7105-04) at 1:2000 in PBS followed by color development with the substrate solution pNPP (Sigma-Millipore, #P7998). After incubating the plates at 37°C for 15 min, the absorbance was read at 405 nm in a plate reader. Absorbance values were corrected for no rHwp1HA control and transformed using the formula: (A_405_ X 20)/15 to reflect the change in absorbance over time of the reactions.

### Western blot

Post *C. albicans* infection, SW480, HCT 116, or LoVo cells were washed twice with PBS and lysed in 100 μL of lysis buffer (60 mM HEPES pH 7.4, 150 mM KCl, 5 mM Na_3_EDTA, 5 mM EGTA, 1 mM Na_3_VO_4_, 50 mM NaF, 1% Triton X-100) (46) with a cocktail of protease inhibitors (Sigma-Aldrich, #P8340) on ice for 15 min. The well contents were scraped and transferred to microfuge tubes and briefly vortexed to ensure complete lysis of the CRC cells. These conditions did not lyse fungal cells. After mixing, the samples were centrifuged (3000 x *g* for 2 min) to pellet fungal cells and cell debris. The supernatants were moved to new tubes and stored at -80°C until gel electrophoresis. Protein concentrations were determined by the Bradford method using a reagent tolerant to detergents (Pierce/ThermoFisher Scientific, #23246). Ten μg of total protein was separated per lane in Tris-glycine 12% acrylamide SDS gels (Novex/Thermo Fisher Scientifc, #XP00125) and transferred to nitrocellulose per standard conditions. The membranes were blocked in 5% BSA, Tris-buffered saline (TBS; 50 mM Tris-HCl, 150 mM NaCl, pH 7.4) for 2 h at room temperature followed by incubation with primary antibodies at 1:1000; rabbit antibodies against E-cadherin (#3195), N-cadherin (#13116), Snail (#3879) (all from Cell Signaling Technology) or 1:10,000 for GAPDH mouse monoclonal antibody (Sigma-Aldrich #G8795) in TBS 0.1% Tween-20 (TBSt) at 4°C for 14-16 h. For detection of MGAT5, membranes were blocked with 5% skim milk in TBSt for 0.5 h followed by incubation with mouse MGAT5 antibody (Biotechne, #MAB5469) at 1:250 in the same buffer at 4°C for 14-16 h. For TG2, membranes were blocked in 5% nonfat dry milk in TBSt for 1 h and incubated in the same buffer with the mouse TG2 antibody used in immunofluorescence assays (ABCAM, #ab2386) at 1:1000 at 4°C for 14-16 h. Washed membranes were incubated with secondary antibodies conjugated to IRDyes (LI-COR) at 1:10,000 dilution in TBSt for 1 h at room temperature protected from light. Washed membranes were scanned (LI-COR Odyssey system) and the band intensities quantified with QuantStudio software (Image Studio Lite v 4.0). Band intensities were quantified from at least 3 independent experiments; in some cases, the data was collected from 6 independent replicates.

For detection of rHwp1HA in ammonium sulfate precipitates, equal volumes from each fraction were subjected to SDS-PAGE and the proteins transferred to an Immobilon P PVDF membrane (Sigma-Aldrich, #IPVH00010) by standard methods. The blot was blocked in 1% skim milk TBSt for 14-16 h at 4°C. The next day, the blot was washed with TBSt and incubated with rabbit anti-HA tag (Thermo Fisher Scientific, # 600-401-384S) at 1:10,000 in blocking solution for 1 h at room temperature. After washing with TBSt, the blot was incubated with goat anti rabbit IgG conjugated to horseradish peroxidase (Thermo Fisher Scientific, #31460) at 1:5000 dilution in block solution for 1 h at room temperature. After washing, HRP activity was detected with a chemiluminescent substrate (Thermo Fisher Scientific, #34580).

### PHA-L lectin binding

SW480, HCT 116, or LoVo cells as near confluent monolayers grown in 12-well plates were infected with WT *C. albican*s at a MOI of 0.5 for 16 h to prepare cell lysates. Other experiments included the MGAT5 inhibitor, swainsonine (Sigma-Aldrich, #S9263) at 1 μg/mL, during the infection. Ten micrograms of total protein was separated by SDS-PAGE and transferred to nitrocellulose. Protein blots were blocked in TBSt for 2 h at room temperature followed by 15 min incubation in 5% BSA in TBSt. The membranes were then incubated with biotinylated PHA-L (Vector Labs, #B-1115-2) at 1:1000 in TBSt for 14-16 h at 4°C. The next day, the blots were washed twice with TBSt and incubated with streptavidin conjugated to IRDye 800CW (LI-COR, #925-32230) at 1:15,000 in TBSt for 1 h protected from light. The blots were washed with TBSt and scanned in a LI-COR Odyssey system. Infections performed in complete media generated a conspicuous protein band of ∼65 kDa reminiscent of BSA absent in control lanes. To avoid over representation of the 65 kDa band in the overall PHA-L binding, protein bands > 65 kDa (SW480) or < 65 kDa (LoVo and HCT 116) were quantified for intensity with QuantStudio software as for western blots.

### N-glycosidase F treatment of cellular glycoproteins

SW480 cells were plated in complete L-15 medium at 2 X 10^5^ in 12-well plates and allowed to reach confluence for 4-5 days. The cell monolayers were washed with HBSS with calcium and magnesium, followed by addition of WT *C. albicans* at an MOI of 0.5 and incubation for 16 h. The monolayers were washed twice with PBS and cell lysates prepared from three replicate cultures on ice as for western blotting. Protein deglycosylation was performed on denatured glycoproteins per the method outlined in (47). Seventy-five μg of total protein in 60 μL with 0.1 % w/v SDS and 100 mM 2-mercaptoethanol (Sigma Aldrich, #M3148) was heated for 5 min at 95°C and cooled on ice. The samples were combined with 8 μL of 10% (w/v) Nonidet P-40 alternative (Millipore/Sigma-Aldrich, #492016), 10 μL of 10X buffer (200 mM Tris–HCl, 250 mM NaCl, 200 mM sodium acetate, pH 7.5), 27 μL molecular grade water, 2 μL of 50X protease inhibitor cocktail (Sigma-Aldrich, #P8340), and 5 U/mL of PNGase F (Sigma-Aldrich, #G1549) to yield a final volume of 100 μL. The samples were incubated for 4 h at 37°C and the reactions terminated by adding 33 μL of 4X reducing Laemmli buffer and heating to 95°C for 5 min. Ten μg of protein was subjected to SDS-PAGE followed by blotting onto nitrocellulose and N-cadherin immunodetection as above. The molecular sizes of N-cadherin species were derived from the migration of prestained Mr protein standards (Novex/Thermo Fisher Scientific, #LC5800).

### Migration assays

Confluent SW480 monolayers in 12-well plates were scratched with a sterile P200 pipet tip in duplicate, washed with PBS to remove floating cells, followed by addition of 625 *C. albicans* SC5314 yeasts added per well to give an MOI of 0.00125 (1 yeast/800 CRC cells) in 0.5 mL of complete media. For HCT 116, or LoVo cells as near confluent monolayers were scratched with a sterile P10 pipet and processed for infection with *C. albicans* in complete media, as above. Multiple areas of each scratch were imaged (Keyence BZ-X710) at the start of the experiment (T0) and at 24 h (T24). Open areas were measured with Photoshop (Adobe; v10.0) and used to calculate percent of wound healing with the formula: 1-(area T24/T0) X 100. The experiment was performed twice with three monolayers per condition in duplicate (n=12). In other experiments, wound healing was measured in the presence of the MGAT5 inhibitor, swainsonine at 1 μg/mL. In these experiments, scratched CRC cell monolayers were briefly washed with PBS, 625 yeasts added (MOI 0.00125), and the cells incubated in complete media with added vehicle (0.1% methanol) or swainsonine for 24 h. Area measurements were taken from three to six independent monolayers per condition in duplicate.

### Candidalysin exposure experiments

The candidalysin peptide [NH_2_]SIIGIIMGILGNIPQVIQIIMSIVKAFKGNK [COOH] (14) (>95% purity; Thermo Fisher Scientific or Peptide Synthetics, Peptide Protein Research, Ltd., Hampshire, UK) was suspended in sterile dH_2_O at 10 mg/mL, aliquoted and stored at -20°C until use. SW480 monolayers (∼5 x 10^5^ cells) were exposed to concentrations of candidalysin (3-70 μM) known to induce a range of epithelial cell responses (14, 48, 49) in complete medium for 2 h. To determine whether candidalysin exposure activated TG2, candidalysin-exposed SW480 cells were washed with HBSS without calcium and magnesium followed by *in situ* TG2 activity assay and detection of crosslinked 5-BPA, as above. In other experiments, 10 μM candidalysin was used to determine the effect of the peptide on E-cadherin, N-cadherin, and Snail1 by western blotting.

### Gene expression assays

Total RNA was prepared from SW480 monolayers infected with WT *C. albicans* (MOI 0.1) for 16 h using a PureLink RNA Mini kit (Invitrogen, #1218301A) according to the manufacturer’s protocol. Contaminating DNA was removed by on-column DNase treatment. Quantitative reverse transcriptase-PCR (qRT-PCR) was performed using SuperScript III SYBR Green One-Step qRT-PCR reagents (Invitrogen, # 12574026), 120-150 ng of total RNA and 10 μM of primers (Integrated DNA Technologies, Coralville, IA) for *CHD1* (For 5’-GCCTCCTGAAAAGAGAGTGGAAG-3’ and Rev 5’-TGGCAGTGTCTCTCCAAATCCG-3’), *OCLN* (For 5’-ATGGCAAAGTGAATGACAAGCGG-3’ and Rev 5’-CTGTAACGAGGCTGCCTGAAGT), *CDLN* (For 5’-GTCTTTGACTCCTTGCTGAATCTG-3’ and Rev 5’-CACCTCATCGTCTTCCAAGCAC-3’), *FN1* (For 5’-ACAACACCGAGGTGACTGAGAC-3’ and Rev 5’-GGACACAACGATGCTTCCTGAG-3’), *SNAI1* (For 5’-TGCCCTCAAGATGCACATCCGA-3’ and Rev 5’-GGGACAGGAGAAGGGCTTCTC-3’) and *GAPDH* (IDT Ready Made primers: For 5’-ACCACAGTCCATGCCATCAC-3’ and Rev 5’-TCCACCACCCTGTTGCTGTA-3’). Relative expression levels were calculated using the ΔΔCt method. Transcript levels were measured from three independent SW480 cultures in triplicate.

### Statistical methods

Data analyses were performed using Prism (GraphPad v10.4.1). P values were calculated using the unpaired Student’s t test or Ordinary one-way ANOVA with significance set a P<0.05. The correlation between 5-BPA+ and rHwp1HA+ cells was quantified using Pearson’s correlation coefficient. P values: *<0.05, **<0.01, ***<0.001; ****<0.0001.

## Results

### *C. albicans* invasion of SW480 CRC cells activates surface TG2

Our previous work identified the *C. albicans* hyphal surface protein, Hwp1, as a substrate for mammalian transglutaminases (12, 42). Extracellular transglutaminase 2 (TG2) is normally inactive, but tissue injury stimulates catalytic activity of intestinal cells *in vivo* and *in vitro* (50). For our studies, we chose CRC model cell lines SW480, HCT 116, and LoVo that differentially express TG2 (i.e., SW480 = high; HCT116 and LoVo = low; See https://depmap.org/portal). TG2, which is uniformly expressed in SW480 cells (51), was activated by crosslinking the TG substrate, 5-(biotinamido)pentylamine (5-BPA) (52, 53) to the cell surface in response to a 3 h infection with wild type (WT) *C. albicans* SC5314 (Fig 1a). 5-BPA+ cells were invaded by *C. albicans* hyphal cells. In contrast, hyphal cells or yeasts bound to the surface of SW480 cells did not activate TG2. Uninfected cells did not display activated surface TG2 activity (Fig. 1b). SW480 cells injured by scratching with a pipet tip also activated TG2 (Fig. 1c) (50). While some *C. albicans*-invaded SW480 cells were negative for 5-BPA suggesting a failure to activate TG2 (Fig. 1d arrows), TG2 was detected by immunofluorescence and immunoblotting (Fig. S1a and b). Cells with activated TG2 (5-BPA-positive) can exhibit decreased immunoreactivity due to conformational changes of the activated enzyme (54). We validated TG2 activity in the crosslinking assays with the TG2-specific inhibitor Z-DON (55) that mimics the glutamine donor substrate. Treatment of SW480 cells with Z-DON (100 μM) abrogated crosslinking of 5-BPA regardless of how the cells were injured (Fig. 1e and f).

**Figure 1.**
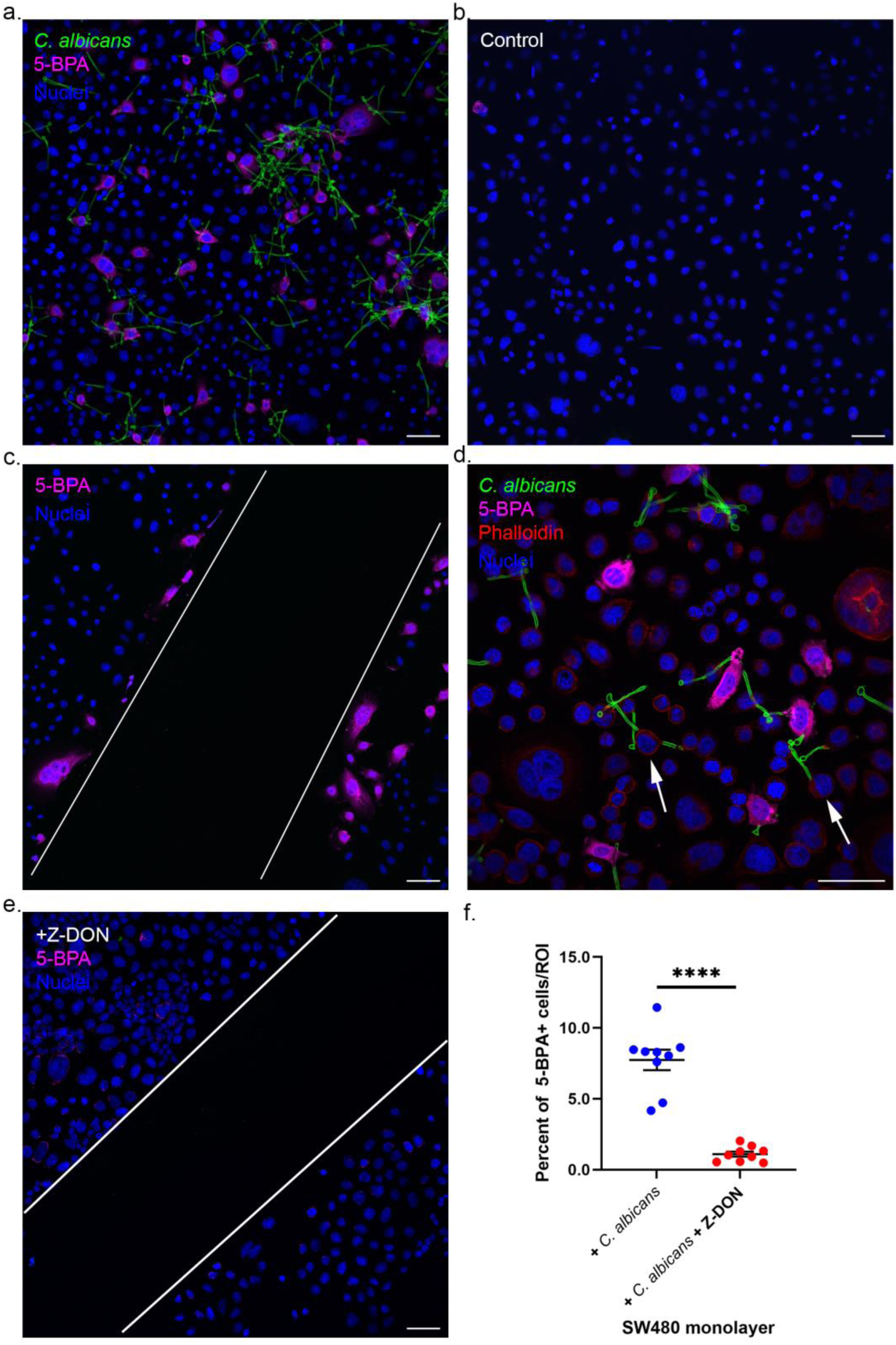
Activation of SW480 TG2 catalytic activity by *C. albicans* cell invasion. SW480 monolayers were infected with *C. albicans* yeasts at an MOI of 2 for 3 h followed by crosslinking 5-BPA for 1 h and fixation. *C. albicans* fungal surfaces were detected with antibodies (green) and 5-BPA with streptavidin-AF647 (magenta). (a) SW480 cells infected with WT *C. albicans* activated cellular TG2 and incorporated the transglutaminase substrate, 5-BPA. (b) Uninfected SW480 cells had few to no cells with highly active TG2. (c) Injuring an SW480 monolayer with a pipet tip induced TG2 activation of wounded cells. (d) A subset of invaded SW480 cells failed to activate TG2 (arrows). (e) The specificity of the 5-BPA crosslinking reaction was validated with the TG2 inhibitor, Z-DON (100 μM). (f) The mean number of 5-BPA positive SW480 cells was significantly diminished when 100 μM Z-DON was included during *C. albicans* infection and in the crosslinking reaction (ROI=9). Significance determined by Student’s t test. **** P<0.0001. Error bars=SEM. Panels a., b., c. and e., magnification 20X. Panel d., magnification 40X. Size bars=50 μm.

In contrast to SW480, *C. albicans* infection of HCT 116 or LoVo cells failed to crosslink 5-BPA (Fig. S1c) consistent with very low TG2 in these cells analyzed by immunoblotting that mirrored published gene expression levels (56) (Fig. S1b). *C. albicans* attached to and readily invaded both HCT 116 and LoVo cells (Fig. S1c arrowheads), suggesting that *C. albicans* adhered to these cells likely utilizing other fungal adhesins, such as Als3 (11, 57–60), independently from covalent attachment through Hwp1 (61–63).

TG2 activity can be affected by Ca^2+^ surges into the cell under cellular stress such as hypoxia or oxidative stress (64). To determine whether *C. albicans* infection induced Ca^2+^ influx, SW480 cells were incubated with the cell permeant intracellular Ca^2+^ chelator, BAPTA-AM (10 μM) and the number of 5-BPA+ cells quantified post *C. albicans* infection (Fig. S1d). While BAPTA-AM treatment appeared to reduce the number of 5-BPA+ cells, these data were not significant (p=0.066). The trend towards fewer 5-BPA-positive cells in BAPTA-AM-treated cells together with published reports (65–67) supports a mechanism of *C. albicans*-induced TG2 activation consistent with Ca^2+^ entry into SW480 cells.

In contrast to CRC cells, *C. albicans* infection of PDO monolayers derived from the colon of healthy subjects (35) did not activate TG2 (Fig. S2). While barrier integrity, as determined by transepithelial electrical resistance (TEER), decreased slightly to ∼86% (+/- 11.5%), the monolayers were resistant to *C. albicans* invasion and did not incorporate 5-BPA. We did not detect TG2 protein PDO cell lysates (Fig. S2b) that was consistent with the lack of detectable TG2 crosslinking activity. Together, the normal human colonic epithelium was resistant to *C. albicans* invasion while CRC cells were permissive to fungal attachment and invasion.

### Candidalysin is sufficient to activate TG2 activity in SW480 cells

Hyphal invasion-mediated epithelial cell injury is critically dependent upon candidalysin, a pore-forming *C. albicans* peptide cytotoxin expressed at the tips of invading hyphae (13, 14, 68) capable of inducing a Ca^2+^ influx into epithelial cells (14). To determine whether cytotoxicity was required to induce TG2 activation, we measured 5-BPA crosslinking to the surface of SW480 cells after infection with a *C. albicans* strain lacking the ability to produce candidalysin (*ece1*Δ/Δ) (Fig. 2a and b). This strain did not activate surface TG2 although it was capable of cell invasion (Fig. 2a) (13, 14, 68). Reintroduction of a functional *ECE1* gene restored TG2 activation, confirming candidalysin as the injury-inducing, TG2-activating factor (Fig 2a and b). The *ece1*Δ/Δ+*ECE1* re-integrant strain did not activate TG2 to the same level as the WT strain (Fig. 2b); however, the increase in the number of 5-BPA+ cells was significantly higher relative to that induced by *ece1*Δ/Δ (Fig.2b). These results suggest that TG2 activation in SW480 cells requires candidalysin.

**Figure 2.**
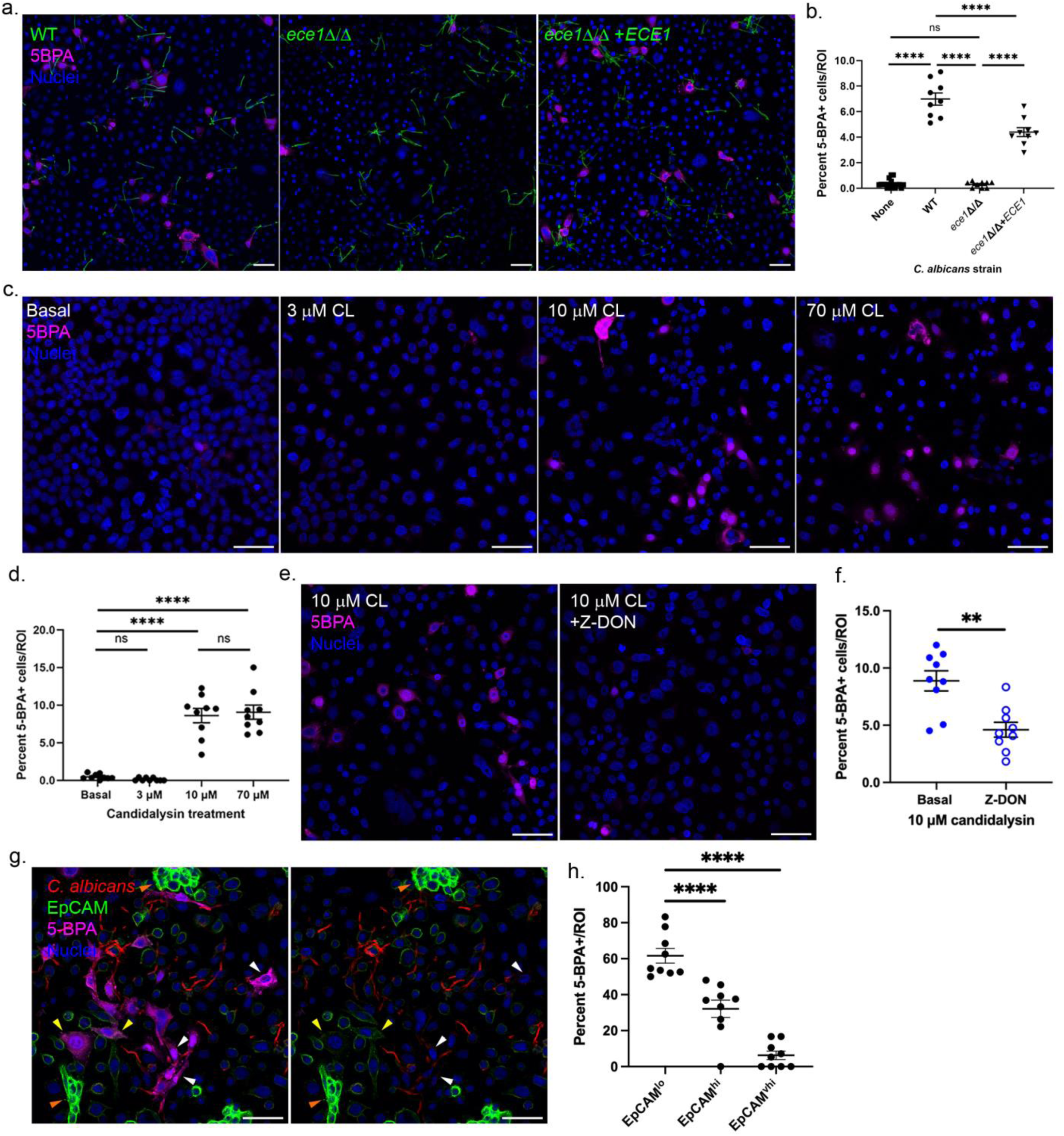
TG2 activation requires candidalysin. SW480 monolayers were infected with *C. albicans* strains (3 h) or treated with candidalysin peptide (CL; 2 h), followed by crosslinking 5-BPA. *C. albicans* fungal surfaces were detected with antibodies (green) and 5-BPA with streptavidin-AF647 (magenta). (a) *C. albicans*-induced TG2 activation is mediated by expression of candidalysin. (b) Mean number of 5-BPA-positve cells post *C. albicans* infection (ROI=9). The candidalysin deletion strain was unable to induce TG2 activation above basal levels. (c) Treatment of SW480 monolayers with candidalysin (CL) peptide activated TG2 in the absence of fungi. (d) Mean number of 5-BPA-positive cells post candidalysin peptide exposure (ROI=9). (e) The specificity of 5-BPA crosslinking catalyzed by TG2 was validated by inhibiting the reaction with 100 μM Z-DON. (f) Z-DON significantly reduced the mean number of 5-BPA-positive cells induced by candidalysin treatment (ROI=9). (g) TG2-activated cells expressed low levels of the epithelial cell marker EpCAM (green; white arrows) detected by immunofluorescence. *C. albicans* surfaces were detected with concanavalin A -TRITC (red). SW480 cells with high expression of EpCAM displayed a spread or spindle morphology (yellow arrows). Cells with very high expression of EpCAM were predominantly negative for 5-BPA crosslinking (orange arrow, bottom left). Significance determined by Ordinary one-way ANOVA in (b) and (d); Significance determined by Student’s t test in (f). **P<0.01; ****P<0.0001. Magnification, 20X. Error bars = SEM. Size bars = 50 μm.

Synthetic candidalysin induces epithelial cell damage in the absence of *Candida* fungal cells (14, 15, 49, 69). Therefore, we asked whether candidalysin alone could activate TG2 on the surface of SW480 cells. Exposure of candidalysin (10 and 70 μM = concentrations that induce epithelial cell signaling responses (48)) to SW480 cells induced TG2 activation (Fig. 2c). The number of 5-BPA+ cells (Fig. 2d) was similar between 10 and 70 μM candidalysin-treated SW480 monolayers, again suggesting that a subset of SW480 cells is susceptible to TG2 activation post injury. Addition of the TG2-specific inhibitor Z-DON confirmed TG2 as the source of crosslinking activity (Fig. 2e and f).

### TG2 unresponsive cells

The SW480 line is a compilation of distinct cell populations of cancer stem cells (70) that may reflect TG2 reactivity. Three distinct SW480 cell subpopulations were recently defined by protein levels of the epithelial cell marker, EpCAM, together with cell morphology: EpCAM^hi^ and EpCAM^lo^ adherent cells, and EpCAM^hi^ spheres (71). We investigated whether TG2-reactivity was associated with these cancer stem cell types based on EpCAM levels and cell morphology. SW480 cells infected with *C. albicans* for 3 h followed by crosslinking 5-BPA were stained for surface expression of EpCAM (Fig. 2g). 5-BPA+ cells were predominately EpCAM-low (gray values ≤30) spindle-like cells (Fig. 2g white arrows). Cells with high EpCAM staining were further stratified into high (EpCAM-high) or very high (EpCAM-vhigh) cells based on plot profile gray values (EpCAM-high 40-60; EpCAM-vhigh >100). EpCAM-high cells positive for 5-BPA tended towards a mesenchymal, spread-out phenotype similar to 5-BPA+/EpCAM-low cells (Fig. 2g yellow arrowheads). TG2 reactivity among EpCAM-vhigh cobblestone-like cells was rare (Fig. 2g and h, orange arrowheads; 6.2% of total 5-BPA+), while 5-BPA+/EpCAM-low cells were the largest population of 5-BPA+ cells (61.6%) even though these cells make up 10-20% of SW480 cells (71) (17.4 +/- 3.7% in our cultures; n=9 ROI). After 3 h of infection, 17.5% (+/- 4.5%; n=17 ROI from two independent experiments) of SW480 cells were invaded by *C. albicans* regardless of EpCAM status. Together, the data suggest the stratification of TG2-activation competent cells as those bearing more mesenchymal characteristics with a low epithelial phenotype. However, not all cells with this phenotype activated TG2 post *C. albicans* invasion.

### *C. albicans*-independent activation of TG2

Candidalysin orthologs are produced by other *Candida* species including the closely related species, *C. tropicalis* (72). We exposed SW480 cells to *C. albicans* (control), *C. tropicalis*, *C. krusei*, *C. auris,* and *C. glabrata* (MOI 2 for 3 h) followed by 5-BPA crosslinking (Fig. S3a and b). We found the wound response to be specific to *C. albicans* as *C. tropicalis* and other pathogenic *Candida* species were unable to activate surface TG2 upon infection of SW480 cells. *C. tropicalis* candidalysin production, like in *C. albicans*, depends upon sustained hyphae cell formation (14, 72) and the poor filamentation of *C. tropicalis* in the SW480 culture likely prevented robust expression of *ECE1* and candidalysin production. Of particular interest was *C. tropicalis* since this species’ fungal DNA together with *C. albicans*’ are positively correlated with late-stage CRC and other gastrointestinal cancers (7, 8). Together, the data showed that TG2 was not activated by other *Candida* spp. secreted virulence factors (e.g., aspartyl proteases) (73, 74) independent of candidalysin.

### TG2 activation and *C. albicans* hyphae adherence

Active TG2 may catalyze tight adherence of new *C. albicans* through Hwp1 expressed on the surface of hyphal elements. Previous work determined that the N-terminal 157 amino acids of Hwp1 contains TG substrate glutamine(s) that participate in crosslinking (12). We re-engineered a recombinant Hwp1 with hemagglutinin (HA) tags at the C-terminus (rHwp1HA; see Methods) to facilitate detection. rHwp1HA was able to crosslink 5-BPA *in vitro* by both human recombinant TG1 or TG2 (Fig. S4a and b). We found that rHwp1HA incorporated onto the surface of cells at the edge of the scratch as observed for 5-BPA (Fig. S4c) suggesting that wounded SW480 cells can bind fungal TG2 substrate, Hwp1.

An important advantage of rHwp1HA is its use as a surrogate for *C. albicans* hyphal cells (12). To determine whether activation of TG2 on the surface of CRC cells promoted crosslinking of new *C. albicans* hyphal cells via Hwp1, SW480 cells were infected with *C. albicans* and followed by crosslinking rHwp1HA. SW480 cells infected with *C. albicans* producing candidalysin (but not *ece1*Δ/Δ) crosslinked rHwp1HA similarly to 5-BPA (Fig. 3a and b) which was dependent on TG2 (Fig. 3c and d). A subpopulation failed to crosslink rHwp1HA even though these were invaded by *C. albicans* hyphae (Fig. 3a arrowheads). We also found that TG2 reactive cells (i.e., rHwp1HA+) expressed low amounts of surface EpCAM (Fig. S4d white arrowheads) similar to 5-BPA+ cells (Fig. 2g). The results showed *C. albicans*-induced activation of surface TG2 via candidalysin produced a functional microenvironment on the surface of SW480 for tight adherence of new *C. albicans* hyphal cells via Hwp1. Together, these data validated rHwp1HA as a substrate for native TG2 on the surface of CRC cells and suggested a molecular mechanism in which *C. albicans* hyphal cells promote their own tight adhesion onto TG2 overexpressing CRC cells.

**Figure 3.**
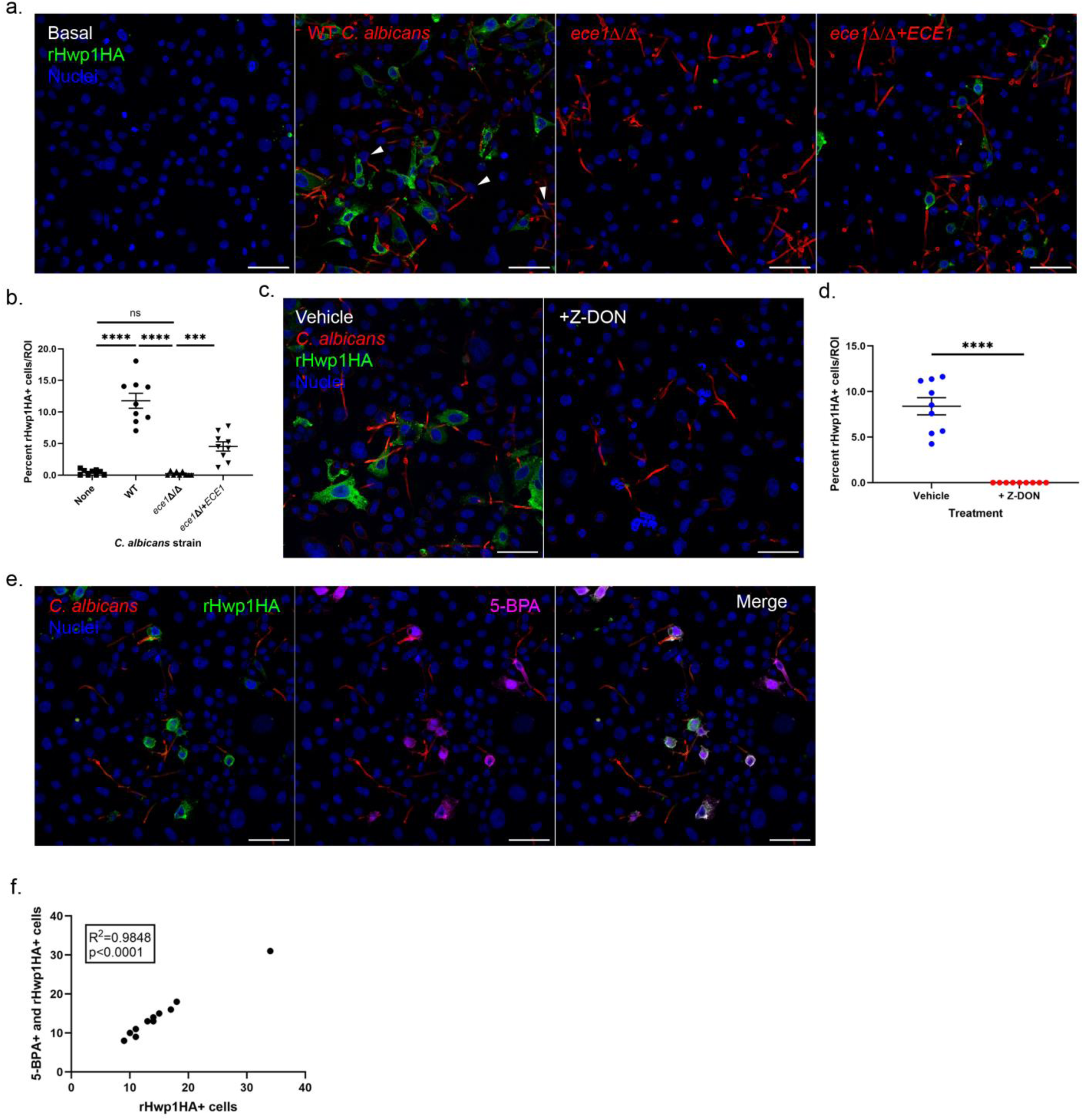
rHwp1 is a substrate for SW480 TG2. SW480 monolayers were infected with *C. albicans* for 3 h followed by rHwp1HA crosslinking. *C. albicans* exposed fungal surfaces were stained with concanavalin-TRITC and antibodies to HA tag were used to detect crosslinked rHwp1HA on cell surfaces. (a and b) TG2 activation by candidalysin crosslinks rHwp1HA. The candidalysin null strain (*ece1*∆/∆) failed to induce rHwp1HA incorporation. Reintroduction of an *ECE1* allele restored TG2 activation (*ece1*∆/∆+*ECE1*). rHwp1HA-positive cells were quantified in 9 ROIs. (c and d) Inhibition of TG2 activity with 100 μM Z-DON prevented rHwp1HA incorporation. (e) Crosslinking of 5-BPA and rHwp1HA occurred on the same cells. 5-BPA was allowed to react first before adding rHwp1HA to the crosslinking reaction. (f) Positive correlation between 5-BPA-positive and rHwpHA-positive cells. (ROI=12). Significance determined by One-way Ordinary ANOVA in (b) and Student’s t test in (d). Covariation was determined by Pearson’s correlation in (f). Magnification 20X. ***P<0.001; ****P<0.0001. Size bars = 50 μm.

Next, we asked whether the TG2-reactive cells crosslinked 5-BPA and rHwp1HA. Because Hwp1 harbors several potential TG-reactive glutamine residues (12, 42), it was important to crosslink 5-BPA (single primary amine) to SW480 cells first to avoid crosslinking 5-BPA directly to rHwp1HA (See Methods for details). Following infection, there was a high degree of colocalization of 5-BPA and rHwp1HA crosslinked to SW480 cells (Fig. 3e and f) suggesting that TG2-reactive cells bear both glutamine (5-BPA crosslinks) and lysine (rHwp1HA crosslinks) TG2 substrate(s), and that these cells make up a single subpopulation of TG2-reactive cells within SW480 cells.

### *C. albicans* invasion of CRC cells affects expression of proteins associated with the epithelial to mesenchymal transition

In addition to modulating TG2 activity, we sought to understand whether *C. albicans* infection of CRC cells affected other aspects of tumor cell physiology. A key event in cancer cell progression occurs via the epithelial to mesenchymal transition (EMT) program. This process is defined by cadherin switching with the loss of the epithelial cell marker, E-cadherin, and the gain of the mesenchymal marker, N-cadherin (75–80). To determine whether *C. albicans* induced the EMT cadherin switch, we probed cell lysates for E-and N-cadherin protein levels prepared from SW480 cells infected with *C. albicans* SC5314 over time (Fig. 4a and b). *C. albicans* infection markedly reduced E-cadherin protein levels at the 16 h time point while N-cadherin resolved as a doublet band with reduced gel mobility reminiscent of posttranslational glycosylation (see below). Based on the maximal EMT-like response of SW480 after 16 h of infection, we chose this time point in subsequent experiments. *C. albicans* infection of HCT 116 and LoVo cells for 16 h produced similar results. E-cadherin levels were reduced while N-cadherin appeared predominantly as a doublet protein band of the same molecular sizes detected in *C. albicans*-infected SW480 cell lysates (Fig. 4c and d). Unlike in SW480 cells, the total amount of N-cadherin in HCT 116 and LoVo cells increased post infection (Fig. 4e). The loss of E-cadherin with concomitant increase in N-cadherin typifies EMT. However, *C. albicans* infection also promoted post-translational modification of N-cadherin in multiple CRC cell lines.

**Figure 4.**
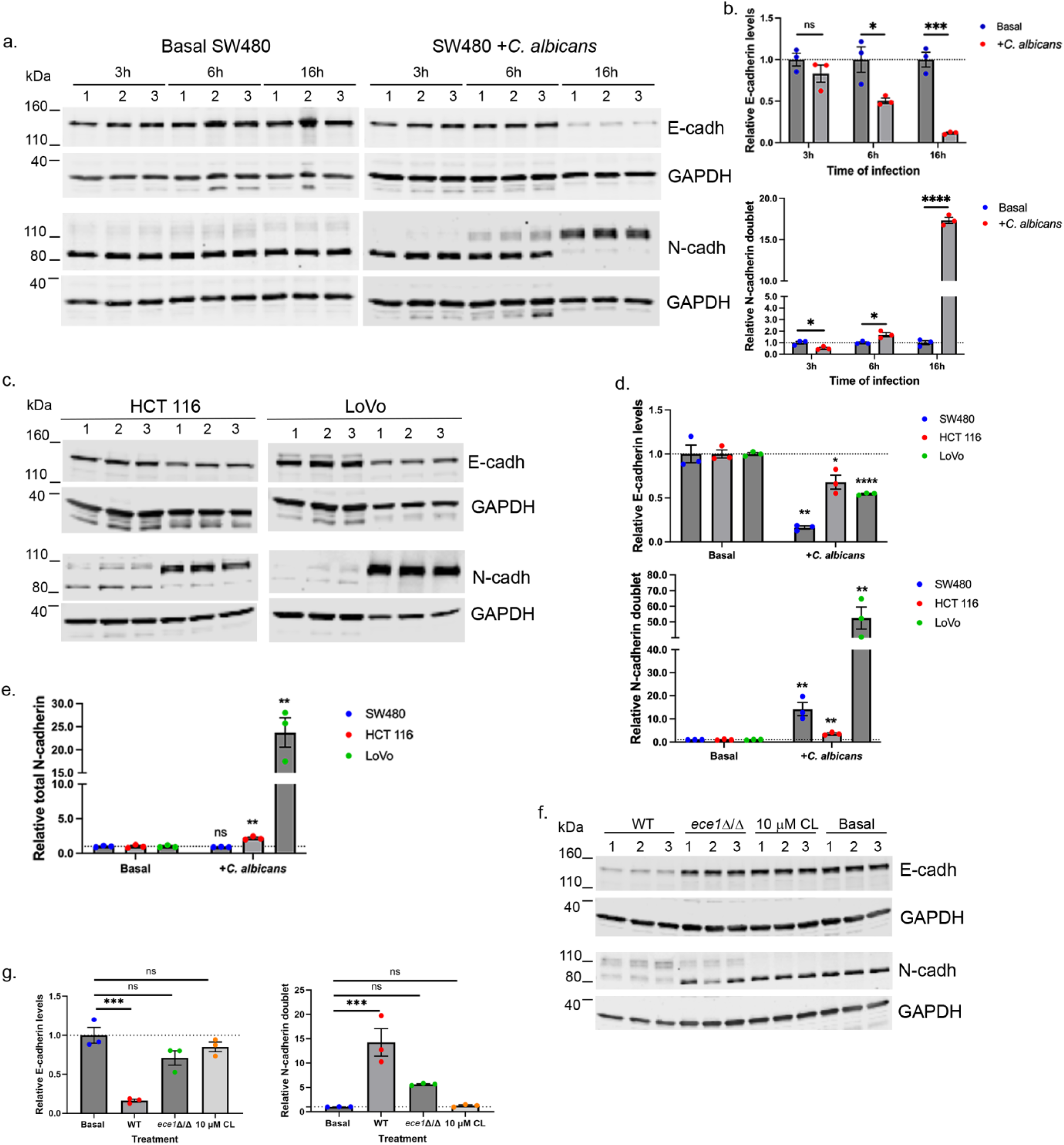
*C. albicans* infection of CRC cells induces EMT-like changes. (a) SW480 cells were infected with WT *C. albicans* over time and SW480 lysates probed for protein levels of E-and N-cadherin by immunoblotting. Three independent infections (n=3; 1, 2, 3) were performed per time point or treatment. (a and b) E-cadherin levels diminished significantly while N-cadherin resolved as a doublet protein as the time of infection was extended. E-cadherin and doublet N-cadherin levels were set relative to uninfected controls (basal). (c) CRC cells HCT 116 and LoVo were infected with WT *C. albicans* for 16 h and cell lysates probed for E-cadherin and N-cadherin. Three independent infections (n=3; 1, 2, 3) were performed per cell line. (d) E-cadherin and doublet N-cadherin levels across all three cell lines. SW480 protein levels from 16 h time point in (b). (e) Total N-cadherin increased in *C. albicans*-infected HCT 116 and LoVo cultures. (f) Production of candidalysin by *C. albicans* augmented reduction of E-cadherin and putative posttranslational modification of N-cadherin (N-cadherin doublet protein bands) in SW480 cells (16 h infections). Candidalysin peptide (CL) had little effect on EMT-like changes. The candidalysin null strain (*ece1*Δ/Δ) induced low level N-cadherin posttranslational modification (ns by Ordinary one-way ANOVA; Student’s t test P=0.0096). Protein levels are set relative to uninfected controls (basal). Significance determined by Ordinary one-way ANOVA, *P<0.05; **P<0.01; ***P<0.001; ****P<0.0001.

Next, we investigated whether candidalysin was necessary to drive EMT-like changes in SW480 cells. SW480 monolayers were infected with *C. albicans* SC5314, *ece1*Δ/Δ or treated with 10 μM candidalysin peptide for 16 h. In cells infected by candidalysin-deficient *C. albicans*, there was a slight reduction in E-cadherin levels, which was similar to cells exposed to synthetic candidalysin (Fig. 4f and g). However, there was a significant increase (P<0.0001) of N-cadherin glycosylation by the *ece1*Δ/Δ strain above control levels although not to the magnitude promoted by WT *C. albicans* (5.64 +/-0.12 vs. 14.24 +/-2.84 fold increase for WT *C. albicans,* P=0.039). Synthetic candidalysin failed to affect glycosylation of N-cadherin suggesting that cell invasion was required for post-translational modification of the protein. These data suggested that *C. albicans* cell invasion coupled with candidalysin toxicity (13) was required for full induction of EMT-type changes, but increased glycosylation of N-cadherin occurred via separate Ca^2+^ independent signaling. Further, exposure of SW480 cells to conditioned media collected from *C. albicans*-infected SW480 monolayers failed to induce any EMT-like changes (Fig. S5a) indicating that active infection was required to see these cellular responses.

Changes in cell differentiation and behavior governed by EMT processes depend upon several transcriptional factors. The upregulation of Snail1 (*SNAI1*) precedes all other EMT-related transcription factors and represses epithelial genes such as *CDH1* (E-cadherin) (81, 82). Both *SNAI1* transcript and protein levels increased in SW480 cells infected with WT *C. albicans* at 16 h relative to controls (Fig. 5a and b). Infection of SW480 cells with *ece1*Δ/Δ strain reduced Snail1 protein levels while exposure with the candidalysin peptide had no effect (Fig. 5c and d; note that the same western blot was used to probe for N-cadherin levels in Fig. 4f). Infection of HCT 116 and LoVo cells with WT *C. albicans* also reduced Snail1 levels in these CRC cells (Fig. 5e and f) similar to SW480 cells infected with the *ece1*Δ/Δ strain (Fig. 5f). However, N-cadherin glycosylation changes did not correlate with Snail1 levels. Further, the decrease of Snail levels in HCT 116 and LoVo cells suggested that these were not sensitive to candidalysin.

**Figure 5.**
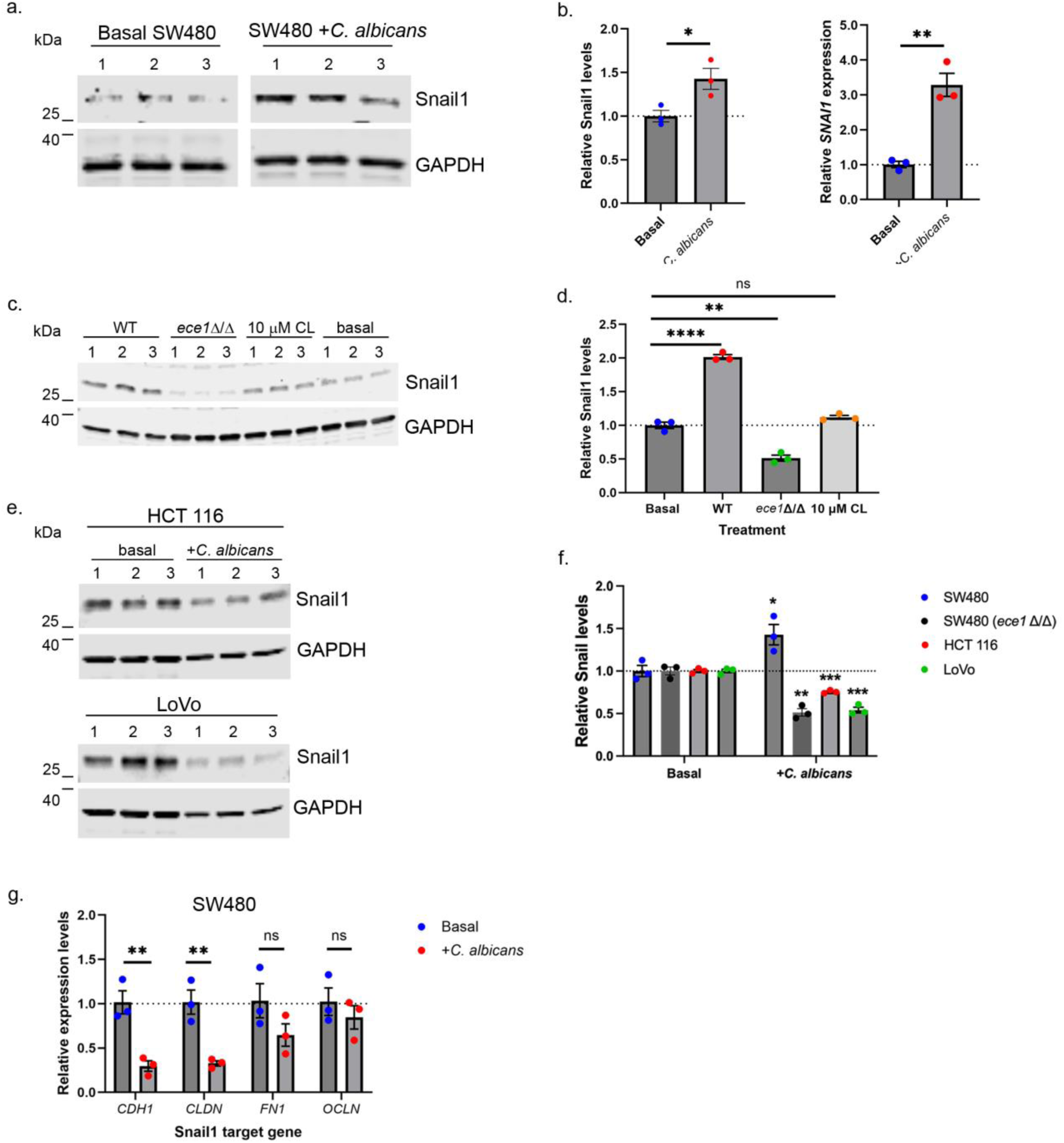
*C. albicans* infection of CRC cells affects expression of the key EMT transcription factor, Snail1. CRC cells were infected with *C. albicans* for 16 h or treated with 10 μM candidalysin (CL) and cell lysates probed for protein levels of Snail1 by immunoblotting. Three independent infections (n=3; 1, 2, 3) were performed per infection or treatment. (a) Increased protein levels of the EMT transcription factor Snail1 post *C. albicans* infection of SW480 cells. The same western blot was analyzed for N-cadherin in Fig. 4f thus GAPDH images are identical. (b) Both Snail1 protein and *SNAI1* transcript abundance increased by 16 h of *C. albicans* infection of SW480 cells. *SNAI1* levels by ΔΔCt relative to uninfected controls (basal). (c and d). Increased Snail1 protein required candidalysin expression by *C. albicans.* The candidalysin null strain (*ece1*Δ/Δ) and candidalysin peptide (CL) failed to upregulate Snail1 protein levels. (e) Infection of HCT 116 and LoVo cells with WT *C. albicans* reduced Snail1 levels similarly to SW480 infected with *ece1*Δ/Δ. (f) Relative amounts of Snail1 among the cell lines; SW480 values from (b) and (c) as comparators. (g) Two out of 4 downstream Snail1 gene targets were affected by increased levels of the transcription factor in SW480 cells. Transcript levels by ΔΔCt relative to uninfected controls (basal). Significance determined by the Student’s t test in (b), (f) and (g). Significance determined by Ordinary one-way ANOVA in (c). *P<0.05; **P<0.01; ***P<0.001; ****P<0.0001.

Surface sulfated glycosaminoglycans (GAGs) mediate candidalysin activity by improving aggregation on the cell surface (38). GAG staining was absent or detected on very few cells in HCT 116 or LoVo cells, respectively. In contrast, SW480 cells stained for GAG at varying intensities across all cells (Fig. S5b). The lack of GAGs may explain the reduced sensitivity of HCT 116 and LoVo cells to candidalysin. Reduced surface expression of GAGs decreased sensitivity to candidalysin toxicity (Fig. S5c). The downregulation of Snail1 paralleled that measured in SW480 cells infected with *ece1*Δ/Δ. The data show that signaling events stemming from the cytotoxic response to WT *C. albicans* infection when candidalysin is delivered in the invasion pocket of GAG-positive SW480 cells upregulated Snail levels and maximized the loss of E-cadherin and glycosylation of N-cadherin (Fig. 5g). The results showed an EMT-like response to *C. albicans* invasion that was augmented when a calcium flux was produced by candidalysin toxicity.

To determine whether the increased abundance of Snail1 affected EMT-related gene targets in SW480 cells, we measured transcript levels of E-cadherin, occludin, claudin 1 (Snail1 repressed) (81–84), and fibronectin (Snail1 upregulated mesenchymal marker) (85, 86). Expression levels of Snail1 targets did not reveal a classical EMT phenotype. While *CHD1* (E-cadherin) and *CLDN* (claudin-1) transcripts were downregulated, *OCLN* (occludin) levels were unchanged, and the expected upregulation of *FN1* (fibronectin) was not observed (Fig. 5g). WT *C. albicans* infection of SW480 cells induced a shift towards a more mesenchymal-like phenotype with canonical loss of epithelial E-cadherin and claudin-1 in the absence of upregulation of fibronectin or increased abundance of N-cadherin. Together, the phenotype of the infected CRC cells was reminiscent of a mixed state, often found within the EMT program that gives rise to a gradation of partial EMT cell states often associated with cancer processes (87).

### *C. albicans* infection affects *N*-glycosylation of N-cadherin and other cellular proteins

Extracellular domains of N-cadherin are *N*-glycosylated with branched β(1–6) *N*-glycans. Increased *N*-glycosylation of N-cadherin reduces cis-dimerization and destabilizes cell to cell adhesion (88), a phenotype associated with increased cell mobility and tumor progression. To determine whether the decreased gel mobility of N-cadherin in *C. albicans*-infected cells was due to increased *N*-glycosylation, SW480 cell lysates were treated with *N*-glycosidase F (PNGase F) and subjected to western blotting (Fig. 6). Removal of *N*-glycans from N-cadherin in *C. albicans*-infected cells increased the protein’s gel migration profile to that of controls (Fig. 6a), confirming that *C. albicans* stimulated posttranslational *N*-glycosylation of N-cadherin. Fungal infection also generated a second de-glycosylated N-cadherin species of reduced molecular size (∼7 kDa smaller; arrow in Fig. 6a) relative to the native protein migrating at 81.5 kDa (N-cadherin has a calculated Mr of 76.7 kDa), suggesting that *C. albicans* additionally induced glycosylation and the expression of a smaller subspecies of N-cadherin.

**Figure 6.**
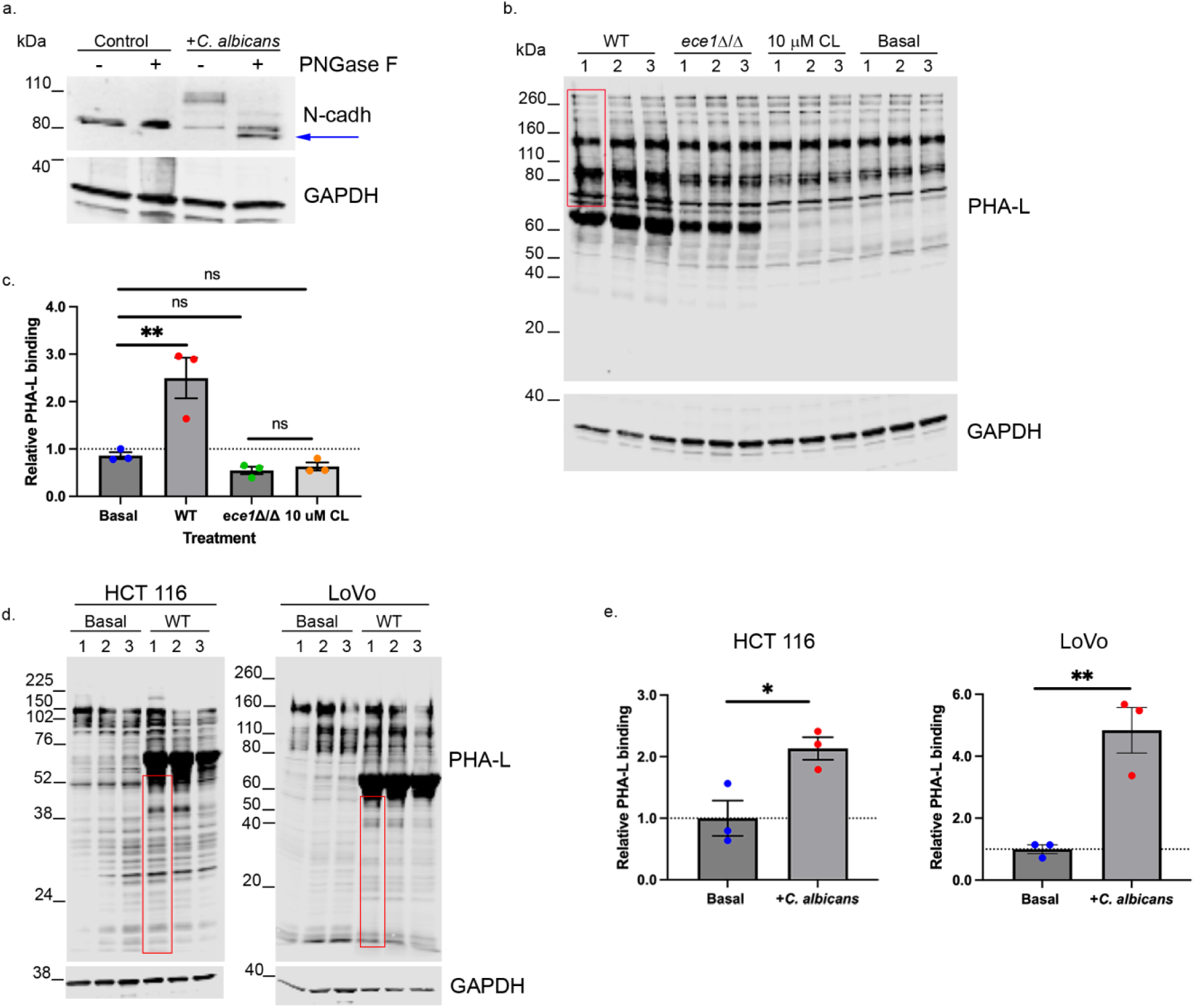
legend. *C. albicans* promotes *N*-glycosylation of N-cadherin and other cellular proteins. (a) Increased N-glycosylation of SW480 post 16 h *C. albicans* infection was validated by treating SW480 lysates with *N*-glycosidase F (PNGase F) followed by western blotting. Deglycosylated N-cadherin resolved as two protein species in lysates from infected SW480 cells; one with native gel mobility and a second faster migrating species (arrow). (b) Global abundance of branched β(1-6) *N*-glycans were probed in SW480 lysates with the lectin PHA-L (*Phaseolus vulgaris* leukoagglutinin) conjugated to biotin. Increased PHA-L binding was quantified in protein species >65k Da (red rectangle) to avoid skewing the data by including putative bovine serum albumin protein band (the growth medium contains FBS). Three independent infections (n=3; 1, 2, 3) were performed per infection or treatment (16 h). PHA-L reactivity was set relative to uninfected controls (basal). (c) The candidalysin null strain (*ece1*Δ/Δ) and candidalysin peptide (CL) did not induce global modification of N-glycans above background. (d and e) *C. albicans* infection (16 h) of HCT 116 and LoVo induced a modest increase in PHA-L reactive proteins of molecular size <65kDa. Infected cultures also harbored a prominent protein band of ∼65kDa (putative BSA from FBS in the culture media). New reactive bands were quantified (red rectangles) as above for SW480 relative to uninfected controls (basal). Three independent infections (n=3; 1, 2, 3) were performed (16 h). Significance determined by Ordinary one-way ANOVA in (c) and Student’s t test in (e). *P<0.05; **P<0.01.

We next asked whether *C. albicans* infection altered global levels of branched β(1–6) N-glycans. To detect these structures we used the lectin PHA-L (*Phaseolus vulgaris* leukoagglutinin), which specifically binds branched β(1–6) *N*-glycans (89), to probe extracts from SW480, HCT 116, and LoVo cells after 16 h *C. albicans* infections. In SW480 extracts, PHA-L bound more strongly to proteins larger than 65 kDa from cells exposed to wild-type *C. albicans* than to those exposed to *ece1*Δ/Δ or to synthetic candidalysin (Fig. 6b and c). In HCT 116 and LoVo lysates, we observed a modest but significant increase in new PHA-L–reactive proteins under 65 kDa in cultures infected with WT *C. albicans* (Fig. 6d and e). Overall, the enhanced PHA-L reactivity in infected cultures indicated that CRC cells mounted a common response to *C. albicans* infection of increased branched *N*-glycosylation.

### *C. albicans* induces cell migration of CRC cells

*N*-glycosylation of N-cadherin destabilizes cell-cell contacts and increases cell migration (88). SW480 cells became more mobile upon exposure to *C. albicans* in a wound-healing (i.e., scratch) assay (Fig. 7a) even at a low multiplicity of infection (MOI 0.00125). Control SW480 cells are not highly migratory; however, exposure to WT *C. albicans* nearly doubled the rate of wound healing in the scratch assay (Fig. 7b). HCT 116 and LoVo cells also showed increased motility when exposed to low concentrations of *C. albicans* (Fig. 7c and d).

**Figure 7.**
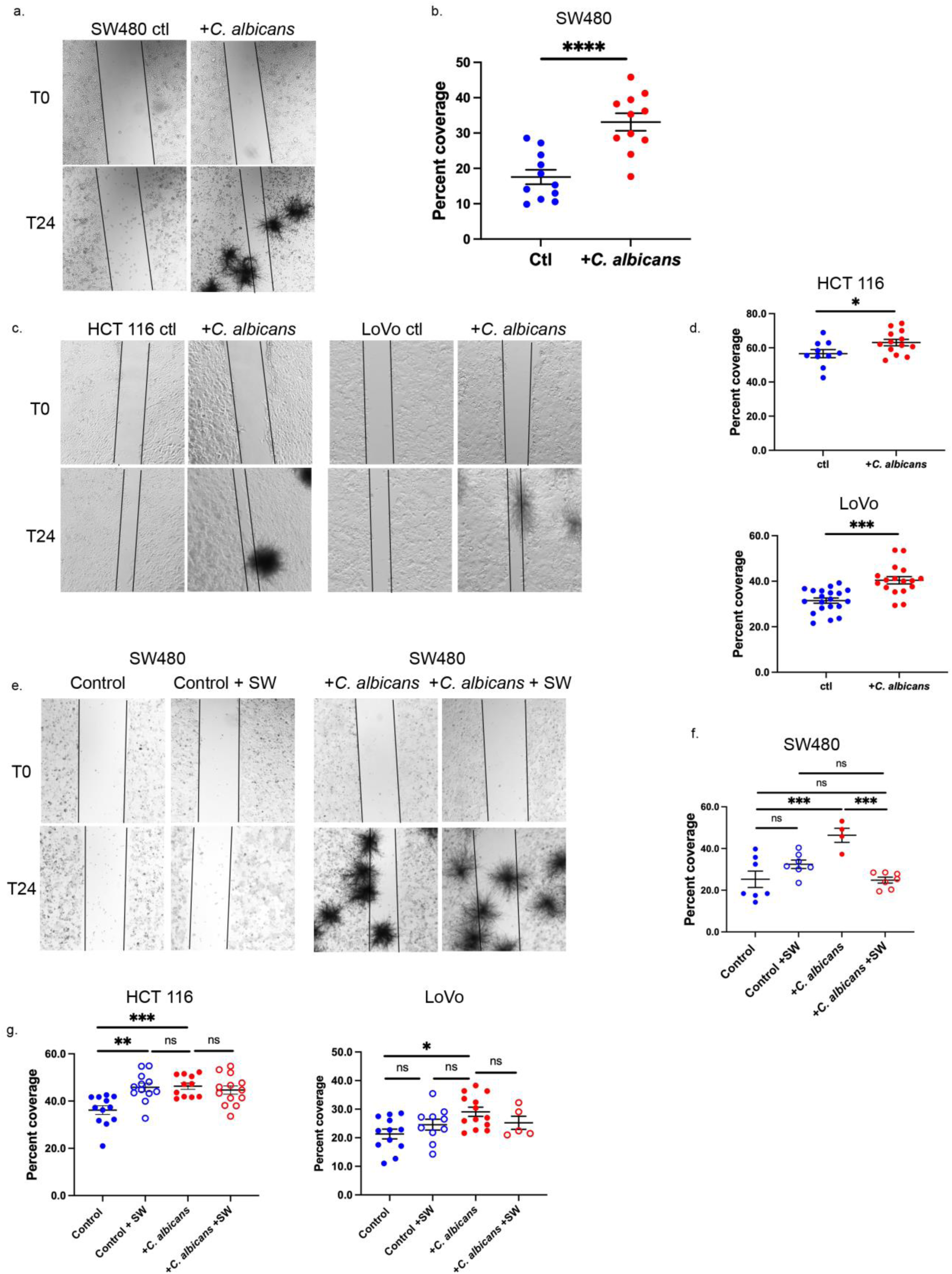
legend. *C. albicans* infection of CRC cells promotes migration. (a and b) *C. albicans* infection of SW480 cells enhanced migration in a wound healing assay. SW480 monolayers were scratched with a pipet tip and allowed to heal for 24 h in the absence or presence of *C. albicans* (MOI 0.00125). Measurements were taken from two independent experiments performed in triplicate (n=11-12). Percent coverage was calculated using the following formula: 1-(area T24/T0) X 100. (c and d) Low numbers of *C. albicans* promote migration of HCT 116 and LoVo cells. Data quantified from three (HCT 116) to six (LoVo) independent monolayers in duplicate (n=10-20). (e and f) Inhibition of β(1-6)-branched *N*-glycosylation with 1 μg/mL of swainsonine (+SW) reduced migration of SW480 cells infected with *C. albicans*. The data was quantified from three independent monolayers in duplicate (n=6-9). (e and f) Inhibition of β(1-6)-branched *N*-glycosylation with 1 μg/mL of swainsonine (+SW) in HCT 116 and LoVo cells did not affect the enhanced migration of *C. albicans*-infected cultures (See representative images in Fig. S6). Data quantified from three independent monolayers in duplicate (n=4-13). Significance in (b and d) was determined by the Student’s t test. Significance in (f) and (g) was determined by Ordinary one-way ANOVA. *P<0.05, **P<0.01, ***P<0.001 and **** P<0.0001. Magnification 4X. Error bars=SEM.

### *C. albicans* infection increases cell migration of SW480 cells and requires branched β(1–6) *N*-<u>glycans activity</u>

The synthesis of branched *N*-glycans recognized by PHA-L is catalyzed by the enzyme N-acetylglucosaminyltransferase V (MGAT5), and increased expression of MGAT5 results in elevated β(1–6)-branched *N*-glycosylation of N-cadherin (88). Based on the increased PHA-L reactivity, we determined whether MGAT5 activity correlated with increased CRC cell migration. To test this, SW480 cells were infected with WT *C. albicans* (MOI 0.00125) in the presence or absence of the alpha-mannosidase II inhibitor swainsonine (90). Swainsonine produces hybrid *N*-glycan structures that lack MGAT5 substrate terminal mannose residues. The migration rate of cells in control monolayers was unaffected by the presence of 1 μg/mL of swainsonine. However, cell migration rates decreased to control levels in swainsonine-treated, scratched SW480 monolayers infected with *C. albicans* (Fig. 7e and f). Swainsonine treatment had little or no effect on the infection-driven increase in migration in HCT 116 and LoVo cells. In HCT 116 cells, swainsonine did not block the enhanced migratory phenotype of infected cultures (Fig. 7g and Fig. S6d) but rather resulted in an unexpected increase in migration at 24 hours post infection. This contrasts with a previous report that 1 μg/mL swainsonine inhibits HCT 116 migration after 48 h (91). Uninfected LoVo cells were insensitive to swainsonine and, like HCT 116, the inhibitor did not reduce the infection-induced enhancement of migration. In contrast, infection of SW480 cells with *C. albicans* did promote migration in a swainsonine-sensitive manner suggesting the increased motility depended upon elevated β(1–6)-branched N-glycosylation. Together, these results suggested that *C. albicans* can influence migratory behavior in CRC cells through upregulation of *N*-glycosylation on N-cadherin and other proteins, although the contribution of this pathway may vary among cell lines.

Because MGAT5 activity contributed to SW480 cell migration in the presence of *C. albicans*, we asked whether MGAT5 activity was related to EMT-like changes induced by fungal infection. Blocking MGAT5 activity did not prevent the loss of E-cadherin, glycosylation of N-cadherin nor the upregulation of Snail1 (Fig. S7a). These data showed the increased MGAT5 activity and EMT-like processes occurred through independent mechanisms that ultimately lead to increased SW480 cell migration in response to *C. albicans* infection.

### *C. albicans* invasion of SW480 CRC cells modulates EMT independently from TG2 activation

Since TG2 activity has been associated with upregulating EMT in CRC and in other cancers (18–22, 92), we speculated that *C. albicans*-induced TG2 activation contributes to EMT progression. To test this, we inhibited TG2 activity in SW480 cells with Z-DON and measured changes in E-cadherin and Snail1 abundance as well as N-cadherin glycosylation following infection with WT *C. albicans* (16 h). HCT 116 and LoVo cells were excluded due to the very low abundance of TG2. Inhibition of TG2 in *C. albicans*-infected SW480 cells did not change E-cadherin levels, prevent *N*-glycosylation of N-cadherin, nor reduce Snail1 protein abundance to control levels (Fig. S7b). TG2 inhibition modestly increased levels of E-cadherin in control cells but did not appreciably affect Snail1 levels or glycosylation of N-cadherin. TG2 inhibition reduced *N*-glycosylation of N-cadherin in SW480 cells infected with *C. albicans* although not to basal levels. TG2 inhibition significantly decreased 5-BPA crosslinking as before (Fig. S7c).

Consistent with the early 3 h infection studies, extending the *C. albicans* infection time to 16 h did not increase the total number of 5-BPA+ cells, although the fungus had overgrown the cell culture growth area and likely invaded the majority of the SW480 cells. Extending the infection time to 16 h decreased the total number of 5-BPA+ cells relative to 3 h (mean 4.45 +/-0.51 vs. 7.74 +/-2.2 cells per ROI, respectively; P=0.004), which may reflect the transitory nature of TG2 activation (50). These data suggest that the EMT changes reflect a cumulative response to *C. albicans* infection as more SW480 cells were invaded over time independent of TG2 activation. The time course experiment (Fig. 4) also supports this cumulative response conclusion. Universal inhibition of TG2 (extracellular and intracellular; Z-DON is cell-permeant) could not rescue the induction of EMT plasticity in SW480 cells together with parallel findings in TG2-low HCT 116 and LoVo cells exposed to *C. albicans* suggesting that the fungus could drive the CRC cells EMT program via a TG2-independent signaling mechanism(s).

## Discussion

Recent reports associate *C. albicans* and colorectal cancer with a poor prognosis (7, 93). An outstanding question is whether *C. albicans* infects tumor tissue because of a favorable growth niche or the presence of the fungus drives tumor evolution. Here we explored both the role of tumor cells in providing a niche for *C. albicans* and the consequences of this interaction on tumor cell differentiation.

The multifunctional enzyme, transglutaminase 2 (TG2), is elevated in CRC (19, 25, 94, 95) and although higher TG2 abundance is correlated with late-stage CRC (25, 95), early-stage CRC tumors have above normal levels of TG2 (25, 94). The human colonic epithelium normally expresses very low mRNA and protein levels of *TGM2* (proteinatlas.org; nTPM <50) (25). In support of these findings, we failed to detect TG2 protein in cell lysates from healthy colon organoids. Infection of PDO monolayers with WT *C. albicans* did not result in cell invasion or incorporation of 5-BPA onto enterocytes after 3 h (Fig. S2). Translocation of *C. albicans* from the lumen of the intestine into the blood stream requires a loss of epithelial barrier function (96, 97) supporting a paracellular mechanism of fungal movement as opposed to an epithelial cell-invasive route of fungal transcytosis. An important biophysical difference between cancer cells and normal epithelial cells is the increased membrane fluidity of cancer cells that affects not only cell shape and rigidity but oncogenic signaling and chemosensitivity (98, 99). Normal epithelial cells tightly control membrane fluidity by modulating cell membrane composition (e.g., cholesterol) in coordination with the cytoskeleton to regulate cell shape and cellular responses. It is conceivable that the more fluid CRC cell membrane allows *C. albicans* to invade these cells while normal colonic epithelial cells are too rigid to permit the prolonged cell membrane invagination produced by penetrating *C. albicans* hyphae (100). CRC cell surface ligands such as EGFR and EphA2 (101–103) that participate in fungal attachment may be absent, inactive, or differentially trafficked in normal epithelial cells. Additional physiological factors such as mucin production, cell polarity, extracellular matrix, and tight junction barrier function are additional features that likely affect the inability of *C. albicans* to invade PDO monolayers. Tumor cells lack many of these attributes in addition to displaying altered surface glycosylation, all factors that shape the epithelial microenvironment and influence *C. albicans* attachment and invasion. The historical use of colon carcinoma Caco-2 cells as a model for *C. albicans* studies of adhesion and invasion of epithelial cells (100–103) together with our CRC cell lines data suggests that neoplastic colonic cells regardless of TG2 levels create a favorable adhesive and invasive environment for *C. albicans* over normal epithelial cells.

Our previous studies of *C. albicans* attachment to human oral epithelial cells show fungal adhesion to host cells via *C. albicans* hyphal protein Hwp1 catalyzed by host transglutaminase activity (12). However, we did not address the mechanism of transglutaminase activity of oral epithelial cells. In these binding assays, senescent buccal cells were scraped from the oral cavity and co-incubated with heat-inactivated *C. albicans* hyphal cells. Although the buccal epithelial cells are senescent, they harbor transglutaminase enzymatic activity provided by either transglutaminase 1 (TG1) or transglutaminase 3 (TG3) in stratified epithelia (104, 105). Here we found that when live *C. albicans* hyphal cells attached and invaded TG2-high CRC SW480 cells (51), this induced an injury response resulting in TG2 activation. Yeasts or hyphal cells remaining external to SW480 cells did not activate TG2. TG2 activation required the production of the cytotoxic peptide candidalysin (14, 15), showing that hyphal invasion of CRC cells without toxicity did not evoke an injury response. Extending the time of infection with the candidalysin deletion strain *ece1*Δ/Δ to 16 h did not activate TG2 above background, indicating that even under prolonged exposure to *C. albicans*, cell toxicity was required to activate TG2 in CRC cells.

We found TG2 activation to be specific to *C. albicans* producing candidalysin. The closely related species, *C. tropicalis*, produces candidalysin with similar biophysical and enhanced cytotoxic properties when the synthetic peptide is tested *in vitro*. However, *C. tropicalis ECE1* expression is highly reduced relative to that measured in *C. albicans* attributed to a decreased formation and maintenance of hyphae in the presence of epithelial cells *in vitro* (72). Not all *C. tropicalis* isolates are equally efficient in producing hyphae and damage to epithelial cells (106, 107). Consistent with these findings, we observed the formation of very short or no *C. tropicalis* hyphae after 3 h in co-culture with CRC cells and no TG2 activation. However, growth within tumor tissue may favor a hyphal morphology. Correlate studies did not specifically address *Candida* species intratumor morphology likely due to the low abundance of fungi coupled with low sensitivity and specificity of fungal staining (8, 93), thus the location and growth morphology of *C. tropicalis* (and other *Candida* species) within GI tumors is not yet understood.

*C. albicans*-induced TG2 activation catalyzed the incorporation of recombinant Hwp1, a surrogate for *C. albicans* hyphal cells (12, 42). This supports a molecular feed-forward mechanism whereby invading *C. albicans* filamentous cells promote the covalent attachment of new hyphae to the surface of TG2-high CRC cells. Strong binding may allow *C. albicans* to persist at CRC tumor niches functionalized by TG2 activity.

A subset of *C. albicans*-invaded or candidalysin-treated SW480 cells failed to activate TG2, reflecting the mixed nature of the SW480 cell line. Transglutaminase protein crosslinking activity depends upon specific glutamine or lysine-bearing substrate proteins (61, 62, 108). The lack of TG2 reactivity of *C. albicans*-invaded or candidalysin-treated SW480 cells likely reflects a cell subpopulation(s) that does not express TG2 protein substrates. Most TG2-reactive cells expressed low surface levels of the epithelial marker, EpCAM, suggesting that mesenchymal-like cells displayed surface-exposed TG2 substrates. EpCAM-low CRC cells tend to populate the tumor front (71), a niche associated with EMT and tumor budding (109, 110). Immunohistochemistry staining of TG2 in CRC tumors shows a general distribution of the enzyme throughout the tumor tissue including at tumor edges (23, 25), supporting a spatial source of tumor-associated enzymatic activity for *C. albicans* attachment by crosslinking. We were able to detect TG2 by immunostaining in all SW480 cells showing that TG2 is over-expressed in these cells (51) regardless of EMT status. Our data does not rule out *C. albicans* attachment to CRC cells by other fungal adhesin proteins (9, 10, 58, 59) through TG2-independent binding; however, Hwp1-mediated covalent attachment may be an added mechanism that enhances fungal persistence at TG2-high CRC tumors.

*C. albicans* invasion of CRC cells perturbed the EMT program in a mixed manner. Fungal infection uniformly decreased expression of the epithelial E-cadherin and increased the global abundance of *N*-glycosylation, including that of N-cadherin. Unlike in SW480 cells, HCT 116 and LoVo cells additionally responded by increasing N-cadherin levels canonically seen in EMT cadherin switching. Cytotoxicity induced by candidalysin enhanced the reduction of E-cadherin and upregulated transcription factor Snail1. The inconsistent downstream regulation of Snail1 targets together with downregulation of Snail1 observed in the absence of candidalysin activity while inducing EMT-like phenotypes suggested that *C. albicans* promoted mixed or hybrid cell states found in intermediate EMT (111, 112). *C. albicans* invasion of OSCC or Caco-2 cells is associated with the proteolytic degradation of E-cadherin (17, 113, 114) and loss of tight junction function without changes in *CDH1* gene expression (114). *C. albicans* infected-OSCC xenographs implanted in mouse tongues show canonical EMT changes with loss of E-cadherin and upregulation of *MMP1*, *MMP10* and *COL5A2* (17). The same group found increased migration of OSCC cells when exposed to *C. albicans in vitro*, showing that *C. albicans* can induce EMT changes and cell mobility in other malignant gastrointestinal epithelial cells.

EMT-like transformation induced by other pathogens or pathobionts such as Vaccinia virus, *Fusobacterium periodonticum*, *Fusobacterium nucleatum*, *Porphyromonas gingivalis* and *Klebsiella pneumonia* have been observed (115–118). Thus EMT-like changes may be a shared cancer cell response to infection or exposure to microbial antigens. In addition, and relevant to our findings, Vaccinia virus, *F. nucleatum,* and heat-killed *P. gingivalis* increase the migratory behavior of oral squamous cell carcinoma epithelial cells (118) as does *F. periodonticum* in esophageal squamous cell carcinoma cells. EMT-like changes are critically associated with metastasis and invasion in CRC (119), with loss of E-cadherin being the key molecular event that enables metastasis (120). Our *in vitro* findings are consistent with the induction of EMT and metastatic-like phenotypes in *C. albicans*-infected CRC cells documented with other oral pathobionts; however, these findings need validation in an *in vivo* CRC model to confirm the migratory phenotype.

Same-cell adhesion can be controlled by the glycosylation status of the extracellular domains of E-and N-cadherins (121). We observed increased *N*-glycosylation of N-cadherin and upregulation of *N*-glycan branching enzymatic activity generated by MGAT5. *C. albicans*-induced cell mobility of SW480 cells was abrogated by inhibiting MGAT5 showing that modifying the glycosylation status of SW480 cells was sufficient to impact cell migration. The inability to diminish HCT 116 and LoVo cell migration by inhibiting MGAT5 suggested several conclusions: other types of migration-associated sugar modifications are concurrently induced, glycosylation-independent migration mechanisms are activated, or the extent of branched *N*-glycosylation was below the threshold to drive cell mobility in these cells when infected with *C. albicans*. Aberrant *N*-glycosylation is a prevailing characteristic of cancer cells, including CRC (122–126), with the addition of branched β (1,6)-linked *N*-acetylglucosamine being a common modification (122, 123). Increased branched *N*-glycosylation can alter the tumor microenvironment in profound ways. In addition to changing same cell adhesion, MGAT5-driven *N*-glycosylation can be associated with an immune-evasive tumor microenvironment through the glycosylation of the T cell-modulating protein Programmed Death-Ligand 1 (PD-L1) and increased infiltration of regulatory T cells (Tregs) (127–129). Clinical MGAT5 inhibitors, which are being developed to reduce tumor growth, metastasis, and immune evasion, remain under investigation. Accordingly, the *C. albicans*-induced increase in MGAT5 activity in CRC cells may have multiple translational implications and warrants further study in *in vivo* CRC models. The induction of aberrant *N*-glycosylation of serum proteins by bacterial, viral and fungal pathogens is an emerging field for the understanding and discovery of new markers of sepsis (130–135). Changes in serum *N*-glycome is thought to originate from host-pathogen interactions of immune cells (i.e., macrophages or dendritic cells) although the exact mechanism is unknown (135). Here we report for the first time the gross alteration of the *N*-glycome of non-immune cells in response to fungal infection. *C. albicans* infection of CRC cells also induced an overall increase in *N*-glycosylation of N-cadherin. *C. albicans* infection additionally generated the expression of a truncated N-cadherin that was *N*-glycosylated; however, it is unclear if this protein species retained functionality. The mechanism of increased *N*-glycosylation and modification of CRC cellular proteins is not yet understood; however, this is consistent with altered *N*-glycosylation of patient serum proteins where there is both an increase in glycosylation and modification of *N*-glycans in response to candidemia (135).

Over expression of TG2 is common in CRC (23–25, 51, 95, 136, 137) and the enzyme’s crosslinking activity modulates and promotes EMT (18–20, 22, 138–141). Because of these phenotypes, we initiated these studies based on the hypothesis that *C. albicans* influences EMT in CRC via TG2 activation. However, we found both TG2-dependent and independent effects on the behavior of CRC cells post *C. albicans* interactions. TG2 activity promoted crosslinking of the hyphae surrogate, Hwp1, to the surface of SW480 cells. At the same time, TG2 catalytic activity was not necessary for EMT-like and -typical changes in *C. albicans*-infected CRC cells, thus, our data did not support our original hypothesis. Inhibition of TG2 in SW480 or very low expression of the enzyme in HCT 116 and LoVo cells did not prevent the loss of E-cadherin or the glycosylation of N-cadherin. In TG2-high CRC tumors, TG2 may induce EMT changes over a longer period beyond our assay time points. TG2 activation is transient (50) and we cannot rule out whether the enzyme’s influence on EMT depends upon repeated cell “injuries” and catalytic re-activation to sustain EMT program signaling.

A limitation of our study was the lack of immune and stromal components that contribute to the CRC tumor microenvironment *in vivo*. The outcome of *in vivo C. albicans*–CRC interactions is likely to be shaped by innate and adaptive immune responses. Neutrophils, macrophages, and dendritic cells may restrict fungal burden, alter hyphal growth and reduce candidalysin exposure, thereby limiting the direct CRC-cell responses observed. Conversely, fungal sensing and epithelial damage can amplify IL-1, IL-17, TNF, TGF-β (48, 142, 143) or myeloid-associated inflammatory circuits that could reinforce EMT-like plasticity, alter glycosylation programs, or promote immune evasion (144, 145). Stromal fibroblasts, endothelial cells, and resident microbiota may further modify TG2 activity, extracellular matrix remodeling, and fungal persistence. The results from our study define CRC cell intrinsic responses to *C. albicans*, but do not fully establish how these mechanisms operate within an intact tumor microenvironment. Increased MGAT5-driven branched *N*-glycosylation creates an immune evasive tumor microenvironment based on PD-L1 glycosylation thus potentially limiting the immune cell contribution of CRC cell responses to *C. albicans* infection in an immune cell/CRC co-culture system. However, our findings should be interpreted as a mechanistic framework that will require validation in more physiologically representative *in vivo* models with a full repertoire of immune cells.

Our data support a model in which *C. albicans* adheres to and invades CRC cells independent of TG2 expression or sensitivity to candidalysin, in contrast to normal colonic epithelium. To isolate the direct effects of infection on cancer cells, we studied CRC lines in the absence of immune, fibroblast, and vascular cells, reasoning that infection of epithelial cells is the initiating event that precedes stromal responses. This approach was also relevant to clinical observations that *Candida*-positive CRC tumors often lack strong inflammatory infiltrates (7, 93), suggesting limited recruitment of inflammatory immune cells in those cases. Despite differences in mutational background and behavior, the CRC cell lines we tested showed consistent responses to C*. albicans*: they adopted EMT-like hybrid characteristics, modified their glycosylation status and became more motile. These foundational findings provide a basis for future work to define the mechanisms driving these responses and to identify potential pharmacological targets to prevent CRC cells from gaining phenotypes associated with metastatic cancer.

## List of Abbreviations

CRC: colorectal cancer
TG1, 2 or 3: transglutaminase 1, 2 or 3
EMT: epithelial to mesenchymal transition
Hwp1: Hyphal wall protein 1
rHwp1HA: recombinant Hwp1 with C-terminal hemagglutinin tag
ROI: region of interest
5-BPA: 5-(biotinamido)pentylamine
MOI: multiplicity of infection
YPD: yeast peptone dextrose
Z-DON: 6-diazo-5-oxo-norleucine tetrapeptide, TG2 Inhibitor
HBSS: Hank’s Balanced Salt solution
PBS: phosphate buffered saline
PHA-L: Phaseolus vulgaris leucoagglutinin lectin
CL: synthetic candidalysin peptide
AF488/568/633: Alexa fluor dyes with emission wavelengths at 488, 568 and 633 nm
PFA: paraformaldehyde
SDS-PAGE: sodium dodecyl sulfate polyacrylamide gel electrophoresis
BSA: bovine serum albumin
TBSt: Tris-buffered saline with 0.5% Tween 20
kDa: kilo Daltons
SW: swainsonine MGAT5 inhibitor
qRT-PCR: Quantitative reverse transcriptase polymerase chain reaction
PNGase F: N-glycosidase F
ANOVA: analysis of variance
SEM: standard error of the mean

## Declarations

### Ethics approval and consent to participate

De-identified patient tissues were obtained with consent under the Johns Hopkins University IRB protocol #NA_00038329, and all methods were performed according to approved guidelines and regulations.

## Consent for publication

Not applicable

## Data availability statement

The authors confirm that the data supporting the findings of this study are available within the article [and/or] its supplementary materials. Data not shown was deposited at the figshare archive under DOI: 10.6084/m9.figshare.29590460.

## Competing interests

The authors declare that they have no competing interests.

## Funding

This work was supported in part by NIH funding grant R21DE021972, the Sherlock Hibbs Endowment and the Johns Hopkins School of Medicine Division of Gastroenterology and Hepatology, to JFS and NIH funding grant P01-AI125181 to NCZ. JRN was supported by the Biotechnology and Biological Sciences Research Council (BBSRC: UKRI717).

## Authors’ contributions

Experimental design, conceptualization, investigation, data visualization: JFS. Formal analysis: JFS and NCZ. Resources (generation of *C. albicans* candidalysin deletion and reconstituted strains): OAP and JRN. Writing-original draft: JFS. Writing-review and editing: JFS, OAP, JRN and NCZ.

## Acknowledgements

We thank Kausik Datta for rHwp1HA analyses and Sean Zhang for providing *Candida* clinical isolates from the Medical Microbiology laboratory at Johns Hopkins Hospital.

---

**Figure S1.** legend. TG2 activity of CRC cells. (a) SW480 monolayer infected with WT *C. albicans* (MOI 2) for 6 h followed by 5-BPA crosslinking for 1 h and fixation. TG2 protein was detected by immunofluorescence (green), *C. albicans* extracellular surfaces with concanavalin A-TRITC (red) and incorporated 5-BPA with streptavidin-AF647 (magenta). Arrows indicate *C. albicans* invaded 5-BPA negative cells expressing TG2. Activated TG2 undergoes a conformational change and becomes less immunoreactive. Magnification= 20X. Size bars = 50 μm. (b) Ten micrograms of total protein was analyzed for the presence of TG2 by immunoblotting. GAPDH is shown as loading control. Table shows the expression level (log-transformed Transcripts Per Million values, TPM) of theTG2 gene (*TGM2*) per line. Expression data sourced from the DepMap portal (https://depmap.org/portal; the Cancer Cell Line Encyclopedia. Release 25Q3). (c) HCT 116 and LoVo monolayers were infected with WT *C. albicans* (MOI 2 for 3 h) followed by 5-BPA crosslinking reactions. Cell surface EpCAM (green) and *C. albicans* extracellular surfaces (red) were detected by immunofluorescence and TG2 activity (incorporated 5-BPA) with streptavidin-AF647 (magenta). Arrows point to examples of *C. albicans*-invaded cells. No 5-BPA-positive cells were observed in multiple ROIs. (d) SW480 cells were infected with *C. albicans* (MOI 2 for 3 h) followed by 5-BPA crosslinking reactions in the absence or presence of the intracellular Ca^2+^ chelator, BAPTA-AM (10 μM). Measurements were taken from 9 ROIs. Magnification = 20X. Size bars = 50 μm.

**Figure S2.** legend. Patient-derived colon organoids are resistant to invasion by WT *C. albicans*. (a) Patient-derived colon organoid (PDO) monolayers were infected with *C. albicans* (MOI 2) for 3 h followed by 5-BPA crosslinking and fixation. The monolayer was permeabilized prior to incubation with antibodies to *C. albicans* (green), phalloidin (actin-red) and streptavidin-AF647 (5-BPA-magenta). *C. albicans* hyphal elements were not observed invading into the monolayer. 5-BPA staining was limited to extracellular material. Top panel, XY projection; bottom panel, XZ projection centered at the yellow cross hairs in the top panel. Magnification 40X. Size bar = 10 μm. (b) Immunodetection of TG2 in PDOs. Two human PDO lines were analyzed for TG2 levels under propagation growth conditions (non-differentiation; ND) or differentiation conditions (DF) for 5 days to induce the developmental differentiation of progenitors into mature colonocytes found at top of colonic crypts (conditions in a. for Colon_1). Ten micrograms of cell lysates were analyzed per lane. GAPDH is shown as loading control.

**Figure S3.** legend. Non-*albicans Candida* fail to activate TG2 activity. (a) SW480 monolayers were infected with *Candida* at an MOI of 2 for 3 h followed by 5-BPA crosslinking and fixation. Extracellular *Candida* surfaces were detected by antibodies (green) and incorporated 5-BPA with streptavidin-AF647 (magenta). Few *C. glabrata* yeasts remained attached to SW480 cells at the end of the assay post multiple washes. (b) The mean number of 5-BPA positive cells was quantified from 9 ROI. Significance was determined by Ordinary one-way ANOVA. Error bars = SEM. Magnification = 20X. Size bars = 50 μm.

**Figure S4.** legend. Production of rHwp1HA and TG substrate activity. (a) Western blot of *Pichia pastoris* culture supernatant expressing rHwp1HA. The proteins in the culture supernatant were precipitated using step-wise concentrations of ammonium sulfate. The precipitated proteins from each fraction where subjected to SDS-PAGE and the proteins transferred to a PVDF membrane. Recombinant Hwp1HA in each fraction was detected with an antibody to the HA tag. Lane (1) starting culture supernatant; (2) 0-25% ammonium sulfate fraction; (3) 25-50% ammonium sulfate fraction enriched for rHwp1HA; (4) 50-77% ammonium sulfate fraction; (5) >77% ammonium sulfate fraction. The fraction in lane (3) (arrow) with the highest amount of rHwp1HA was desalted by dialysis, concentrated and used as source of rHwp1HA. No other protein was detected in the concentrated sample by Coomassie blue staining of a separate SDS-PAG. The migration of protein M_r_ standards is shown at left. (b) In vitro transglutaminase assays using increasing concentrations of rHwp1HA, 50 μM 5-BPA and 1 U/mL of either human transglutaminase 1 (TG1) or human transglutaminase 2 (TG2). 5-BPA crosslinked to rHwp1HA was determined by the change in absorbance over 15 min. The data are means of two independent experiments performed in triplicate. (c) rHwp1HA serves as a substrate for native TG2 on the surface of SW480 cells. An SW480 monolayer was scratched and incubated in TG2 reaction buffer with 2.5 μM rHwp1HA for 1 h at 37°C. Crosslinked rHwp1HA was detected with antibodies to the HA tag (green). Magnification = 20X. Size bars = 50 μm.

**Figure S5.** legend. Conditioned media does not induce EMT-like changes. HCT 116 and LoVo cells do not express cell-surface sulfated glycosaminoglycans (GAGs). (a) SW480 monolayers were exposed to conditioned media from SW480 or SW480 *C. albicans*-infected cultures (MOI 0.1 for 24 h) for 16 h and E-cadherin and N-cadherin levels analyzed by immuoblotting. (b) Cell surface GAGs (green) were detected by immunofluorescence in uninfected cultures. Magnification = 20X. Size bars = 50 μm. (c) Candidalysin cytotoxicity was measured in GAG-positive (SW480) and negative (LoVo) cells by measuring release of lactate dehydrogenase (LDH). Triplicate wells were seeded with 20,000 cells for 48 h and the wells treated with candidalysin for 24 h. Percent toxicity was determined from total LDH release induced by incubating control wells in 10% tergitol. Significance by Student’s t Test. ****P<0.0001

**Figure S6.** **legend. Inhibition of branched β(1-6) *N*-glycosylation does not prevent *C.*** *albicans*-enhanced cell migration of HCT 116 and LoVo cells. HCT 116 and LoVo near confluent monolayers were scratched with a 10 μL pipette tip and infected with WT *C. albicans* (MOI 0.00125) for 24 h. Scratched images were captured at the start (T0) and at 24 h (T24). Addition of the MGAT5 activity inhibitor, swainsonine (+SW) at 1μg/mL had no effect on *C. albicans*-enhanced cell migration. Representative images are shown. Magnification = 4X.

**Figure S7.** legend. EMT-like changes were induced by *C. albicans* infection when branched β(1-6) *N*-glycosylation and TG2 activity were inhibited. Decreased TG2 activity after 16 h *C. albicans* infection. (a) Immunoblot analysis of SW480 lysates from cells infected with WT *C. albicans* for 16 h (MOI 0.5) when branched *N*-glycosylation was inhibited with 1 μg/mL swainsonine (+SW). *C. albicans*-induced EMT-like changes were not affected by inhibiting branched *N*-glycosylation. (b) Immunoblot analyses of SW480 lysates prepared from cells infected with WT *C. albicans* for 16 h (MOI 0.5) in the presence (+Z-DON) or absence (Vehicle) of the TG2 inhibitor, Z-DON (100 μM). The cell extracts were probed for levels of E-cadherin, glycosylated N-cadherin and abundance of the EMT transcription factor, Snail1. TG2 inhibition did not prevent EMT-like changes in infected cultures. M_r_ markers are indicated at left. The same western blot was probed for N-cadherin and Snail1. (c) SW480 infected with WT *C. albicans* for 16 h followed by a TG2 activity assay (5-BPA incorporation). *C. albicans* surfaces were detected by immunofluorescence (green) and TG2 active cells (magenta) with streptavidin-647. The number of 5-BPA positive cells was quantified from 9 ROIs. Magnification = 20X. Size bars = 50 μm. Significance in (a) and (b) determined by Ordinary one-way ANOVA and by the Student’s t Test in (c). **P<0.01; ***P<0.001; ****P<0.0001.

**Supplementary Figure 8.** Uncropped western blot images. Source data for figures 4, 5, 6, S1, S2, S5 and S7.

## Notes

### Competing Interest Statement

The authors have declared no competing interest.

## References

1. Jasperson KW, Tuohy TM, Neklason DW, Burt RW. Hereditary and familial colon cancer. Gastroenterology. 2010;138(6):2044–58.

2. Keum N, Giovannucci E. Global burden of colorectal cancer: emerging trends, risk factors and prevention strategies. Nat Rev Gastroenterol Hepatol. 2019;16(12):713–32.

3. Mei S, Deng Z, Chen Y, Ning D, Guo Y, Fan X, et al. Dysbiosis: The first hit for digestive system cancer. Front Physiol. 2022;13:1040991.

4. Niekamp P, Kim CH. Microbial Metabolite Dysbiosis and Colorectal Cancer. Gut Liver. 2023.

5. Sears CL, Garrett WS. Microbes, microbiota, and colon cancer. Cell Host Microbe. 2014;15(3):317–28.

6. Zafari N, Velayati M, Mehrabadi S, Damavandi S, Khazaei M, Hassanian SM, et al. Remodeling of the Gut Microbiota in Colorectal Cancer and its Association with Obesity. Curr Pharm Des. 2023.

7. Dohlman AB, Klug J, Mesko M, Gao IH, Lipkin SM, Shen X, et al. A pan-cancer mycobiome analysis reveals fungal involvement in gastrointestinal and lung tumors. Cell. 2022;185(20):3807–22 e12.

8. Oba S, Okuno K, Watanabe S, Yamamoto Y, Takaoka A, Hanaoka M, et al. Intratumoral fungal burden of Candida tropicalis as a novel prognostic biomarker for recurrence and mortality in colorectal cancer. Cancer. 2026;132(8):e70408.

9. Hiller E, Zavrel M, Hauser N, Sohn K, Burger-Kentischer A, Lemuth K, et al. Adaptation, adhesion and invasion during interaction of Candida albicans with the host--focus on the function of cell wall proteins. Int J Med Microbiol. 2011;301(5):384–9.

10. Hostetter MK. Linkage of adhesion, morphogenesis, and virulence in Candida albicans. J Lab Clin Med. 1998;132(4):258–63.

11. Liu Y, Filler SG. Candida albicans Als3, a multifunctional adhesin and invasin. Eukaryot Cell. 2011;10(2):168–73.

12. Staab JF, Bradway SD, Fidel PL, Sundstrom P. Adhesive and mammalian transglutaminase substrate properties of Candida albicans Hwp1. Science. 1999;283(5407):1535–8.

13. Mogavero S, Sauer FM, Brunke S, Allert S, Schulz D, Wisgott S, et al. Candidalysin delivery to the invasion pocket is critical for host epithelial damage induced by Candida albicans. Cell Microbiol. 2021;23(10):e13378.

14. Moyes DL, Wilson D, Richardson JP, Mogavero S, Tang SX, Wernecke J, et al. Candidalysin is a fungal peptide toxin critical for mucosal infection. Nature. 2016;532(7597):64–8.

15. Nikou SA, Zhou C, Griffiths JS, Kotowicz NK, Coleman BM, Green MJ, et al. The Candida albicans toxin candidalysin mediates distinct epithelial inflammatory responses through p38 and EGFR-ERK pathways. Sci Signal. 2022;15(728):eabj6915.

16. Dickenson RE, Pellon A, Ponde NO, Hepworth O, Daniels Gatward LF, Naglik JR, et al. EGR1 regulates oral epithelial cell responses to Candida albicans via the EGFR-ERK1/2 pathway. bioRxiv. 2023.

17. Vadovics M, Ho J, Igaz N, Alfoldi R, Rakk D, Veres E, et al. Candida albicans Enhances the Progression of Oral Squamous Cell Carcinoma In Vitro and In Vivo. mBio. 2021;13(1):e0314421.

18. Ayinde O, Wang Z, Griffin M. Tissue transglutaminase induces Epithelial-Mesenchymal-Transition and the acquisition of stem cell like characteristics in colorectal cancer cells. Oncotarget. 2017;8(12):20025–41.

19. Ayinde O, Wang Z, Pinton G, Moro L, Griffin M. Transglutaminase 2 maintains a colorectal cancer stem phenotype by regulating epithelial-mesenchymal transition. Oncotarget. 2019;10(44):4556–69.

20. Blaheta RA, Han J, Oppermann E, Bechstein WO, Burkhard K, Haferkamp A, et al. Transglutaminase 2 promotes epithelial-to-mesenchymal transition by regulating the expression of matrix metalloproteinase 7 in colorectal cancer cells via the MEK/ERK signaling pathway. Biochim Biophys Acta Mol Basis Dis. 2025;1871(1):167538.

21. Cao L, Shao M, Schilder J, Guise T, Mohammad KS, Matei D. Tissue transglutaminase links TGF-beta, epithelial to mesenchymal transition and a stem cell phenotype in ovarian cancer. Oncogene. 2012;31(20):2521–34.

22. Lin CY, Tsai PH, Kandaswami CC, Chang GD, Cheng CH, Huang CJ, et al. Role of tissue transglutaminase 2 in the acquisition of a mesenchymal-like phenotype in highly invasive A431 tumor cells. Mol Cancer. 2011;10:87.

23. Fernandez-Acenero MJ, Torres S, Garcia-Palmero I, Diaz Del Arco C, Casal JI. Prognostic role of tissue transglutaminase 2 in colon carcinoma. Virchows Arch. 2016;469(6):611–9.

24. Malkomes P, Lunger I, Oppermann E, Abou-El-Ardat K, Oellerich T, Gunther S, et al. Transglutaminase 2 promotes tumorigenicity of colon cancer cells by inactivation of the tumor suppressor p53. Oncogene. 2021;40(25):4352–67.

25. Malkomes P, Lunger I, Oppermann E, Lorenz J, Faqar-Uz-Zaman SF, Han J, et al. Transglutaminase 2 is associated with adverse colorectal cancer survival and represents a therapeutic target. Cancer Gene Ther. 2023.

26. Miyoshi N, Ishii H, Mimori K, Tanaka F, Hitora T, Tei M, et al. TGM2 is a novel marker for prognosis and therapeutic target in colorectal cancer. Ann Surg Oncol. 2010;17(4):967–72.

27. Motta C, Pellegrini A, Camaione S, Geoghegan J, Speziale P, Barbieri G, et al. von Willebrand factor-binding protein (vWbp)-activated factor XIII and transglutaminase 2 (TG2) promote cross-linking between FnBPA from Staphylococcus aureus and fibrinogen. Sci Rep. 2023;13(1):11683.

28. Nicoll WS, Sacci JB, Rodolfo C, Di Giacomo G, Piacentini M, Holland ZJ, et al. Plasmodium falciparum liver stage antigen-1 is cross-linked by tissue transglutaminase. Malar J. 2011;10:14.

29. Ponniah G, Rollenhagen C, Bahn YS, Staab JF, Sundstrom P. State of differentiation defines buccal epithelial cell affinity for cross-linking to Candida albicans Hwp1. J Oral Pathol Med. 2007;36(8):456–67.

30. Sundstrom P, Balish E, Allen CM. Essential role of the Candida albicans transglutaminase substrate, hyphal wall protein 1, in lethal oroesophageal candidiasis in immunodeficient mice. J Infect Dis. 2002;185(4):521–30.

31. Gillum AM, Tsay EY, Kirsch DR. Isolation of the Candida albicans gene for orotidine-5’-phosphate decarboxylase by complementation of S. cerevisiae ura3 and E. coli pyrF mutations. Mol Gen Genet. 1984;198(2):179–82.

32. Paulin OKA, Tsavou A, Priest EL, Griffiths JS, Lortal L, Kempf A, et al. The combinatorial action of hyphal growth and candidalysin is critical for promoting Candida albicans oropharyngeal infection. mBio. 2026;17(1):e0330425.

33. Nguyen N, Quail MMF, Hernday AD. An Efficient, Rapid, and Recyclable System for CRISPR-Mediated Genome Editing in Candida albicans. mSphere. 2017;2(2).

34. Rose MW, F.; Hieter, P. Methods in yeast genetics — A laboratory course manual. In: Radford A, editor. 19. Cold Spring Harbor, New York: Cold Spring Harbor Laboratory Press; 1990. p. 198.

35. Noel G, Baetz NW, Staab JF, Donowitz M, Kovbasnjuk O, Pasetti MF, et al. A primary human macrophage-enteroid co-culture model to investigate mucosal gut physiology and host-pathogen interactions. Sci Rep. 2017;7:45270.

36. Staab JF, Lemme-Dumit JM, Latanich R, Pasetti MF, Zachos NC. Co-Culture System of Human Enteroids/Colonoids with Innate Immune Cells. Curr Protoc Immunol. 2020;131(1):e113.

37. Fujii M, Matano M, Nanki K, Sato T. Efficient genetic engineering of human intestinal organoids using electroporation. Nat Protoc. 2015;10(10):1474–85.

38. Lin J, Miao J, Schaefer KG, Russell CM, Pyron RJ, Zhang F, et al. Sulfated glycosaminoglycans are host epithelial cell targets of the Candida albicans toxin candidalysin. Nat Microbiol. 2024;9(10):2553–69.

39. Gulati M, Lohse MB, Ennis CL, Gonzalez RE, Perry AM, Bapat P, et al. In Vitro Culturing and Screening of Candida albicans Biofilms. Curr Protoc Microbiol. 2018;50(1):e60.

40. Richardson MD, Kearns MJ, Smith H. Differentiation of extracellular from ingested Candida albicans blastospores in phagocytosis tests by staining with fluorescein-labelled concanavalin A. J Immunol Methods. 1982;52(2):241–4.

41. Wickramasinghe DN, Lyon CM, Lee S, Hepworth OW, Priest EL, Maufrais C, et al. Variations in candidalysin amino acid sequence influence toxicity and host responses. mBio. 2024;15(8):e0335123.

42. Staab JF, Bahn YS, Tai CH, Cook PF, Sundstrom P. Expression of transglutaminase substrate activity on Candida albicans germ tubes through a coiled, disulfide-bonded N-terminal domain of Hwp1 requires C-terminal glycosylphosphatidylinositol modification. J Biol Chem. 2004;279(39):40737–47.

43. Butterworth PJ. A practical guide to enzymology. Clarence H. Suelter, Wiley Interscience, 281 + xiii pages. Price £46 in hard cover (1985). Cell Biochemistry and Function. 1986;4(4):303-.

44. Scopes RK. Measurement of protein by spectrophotometry at 205 nm. Anal Biochem. 1974;59(1):277–82.

45. Stoscheck CM. Quantitation of protein. Methods Enzymol. 1990;182:50–68.

46. Foulke-Abel J, In J, Yin J, Zachos NC, Kovbasnjuk O, Estes MK, et al. Human Enteroids as a Model of Upper Small Intestinal Ion Transport Physiology and Pathophysiology. Gastroenterology. 2016;150(3):638–49 e8.

47. Sinyuk M, Lathia JD, Viapiano MS. Characterization and Analysis of Extracellular Matrix in Malignant Brain Tumors and Their Cellular Derivatives. In: Leach JB, Powell EM, editors. Extracellular Matrix. New York, NY: Springer New York; 2015. p. 113–38.

48. Ho J, Yang X, Nikou SA, Kichik N, Donkin A, Ponde NO, et al. Candidalysin activates innate epithelial immune responses via epidermal growth factor receptor. Nat Commun. 2019;10(1):2297.

49. Ponde NO, Lortal L, Tsavou A, Hepworth OW, Wickramasinghe DN, Ho J, et al. Receptor-kinase EGFR-MAPK adaptor proteins mediate the epithelial response to Candida albicans via the cytolytic peptide toxin, candidalysin. J Biol Chem. 2022;298(10):102419.

50. Siegel M, Strnad P, Watts RE, Choi K, Jabri B, Omary MB, et al. Extracellular transglutaminase 2 is catalytically inactive, but is transiently activated upon tissue injury. PLoS One. 2008;3(3):e1861.

51. Zirvi KA, Keogh JP, Slomiany A, Slomiany BL. Transglutaminase activity in human colorectal carcinomas of differing metastatic potential. Cancer Lett. 1991;60(1):85–92.

52. Ikura K, Kita K, Fujita I, Hashimoto H, Kawabata N. Identification of amine acceptor protein substrates of transglutaminase in liver extracts: use of 5-(biotinamido) pentylamine as a probe. Arch Biochem Biophys. 1998;356(2):280–6.

53. Singh RN, McQueen T, Mehta K. Detection of the amine acceptor protein substrates of transglutaminase with 5-(biotinamido) pentylamine. Anal Biochem. 1995;231(1):261–3.

54. Kumar A, Hu J, LaVoie HA, Walsh KB, DiPette DJ, Singh US. Conformational changes and translocation of tissue-transglutaminase to the plasma membranes: role in cancer cell migration. BMC Cancer. 2014;14:256.

55. McConoughey SJ, Basso M, Niatsetskaya ZV, Sleiman SF, Smirnova NA, Langley BC, et al. Inhibition of transglutaminase 2 mitigates transcriptional dysregulation in models of Huntington disease. EMBO Mol Med. 2010;2(9):349–70.

56. Arafeh R, Shibue T, Dempster JM, Hahn WC, Vazquez F. The present and future of the Cancer Dependency Map. Nat Rev Cancer. 2025;25(1):59–73.

57. Cota E, Hoyer LL. The Candida albicans agglutinin-like sequence family of adhesins: functional insights gained from structural analysis. Future Microbiol. 2015;10(10):1635–548.

58. Martin H, Kavanagh K, Velasco-Torrijos T. Targeting adhesion in fungal pathogen Candida albicans. Future Med Chem. 2021;13(3):313–34.

59. Rosiana S, Zhang L, Kim GH, Revtovich AV, Uthayakumar D, Sukumaran A, et al. Comprehensive genetic analysis of adhesin proteins and their role in virulence of Candida albicans. Genetics. 2021;217(2).

60. Sundstrom P. Adhesins in Candida albicans. Curr Opin Microbiol. 1999;2(4):353–7.

61. Greenberg CS, Birckbichler PJ, Rice RH. Transglutaminases: multifunctional cross-linking enzymes that stabilize tissues. FASEB J. 1991;5(15):3071–7.

62. Lorand L, Conrad SM. Transglutaminases. Mol Cell Biochem. 1984;58(1-2):9–35.

63. Lorand L, Graham RM. Transglutaminases: crosslinking enzymes with pleiotropic functions. Nat Rev Mol Cell Biol. 2003;4(2):140–56.

64. Kiraly R, Demeny M, Fesus L. Protein transamidation by transglutaminase 2 in cells: a disputed Ca2+-dependent action of a multifunctional protein. FEBS J. 2011;278(24):4717–39.

65. Jeong EM, Kim CW, Cho SY, Jang GY, Shin DM, Jeon JH, et al. Degradation of transglutaminase 2 by calcium-mediated ubiquitination responding to high oxidative stress. FEBS Lett. 2009;583(4):648–54.

66. Jung HJ, Chen Z, Wang M, Fayad L, Romaguera J, Kwak LW, et al. Calcium blockers decrease the bortezomib resistance in mantle cell lymphoma via manipulation of tissue transglutaminase activities. Blood. 2012;119(11):2568–78.

67. Yoo JO, Lim YC, Kim YM, Ha KS. Transglutaminase 2 promotes both caspase-dependent and caspase-independent apoptotic cell death via the calpain/Bax protein signaling pathway. J Biol Chem. 2012;287(18):14377–88.

68. Westman J, Plumb J, Licht A, Yang M, Allert S, Naglik JR, et al. Calcium-dependent ESCRT recruitment and lysosome exocytosis maintain epithelial integrity during Candida albicans invasion. Cell Rep. 2022;38(1):110187.

69. Ho J, Wickramasinghe DN, Nikou SA, Hube B, Richardson JP, Naglik JR. Candidalysin Is a Potent Trigger of Alarmin and Antimicrobial Peptide Release in Epithelial Cells. Cells. 2020;9(3).

70. Brabletz T, Jung A, Spaderna S, Hlubek F, Kirchner T. Opinion: migrating cancer stem cells - an integrated concept of malignant tumour progression. Nat Rev Cancer. 2005;5(9):744–9.

71. Verhagen MP, Xu T, Stabile R, Joosten R, Tucci FA, van Royen M, et al. The SW480 cell line as a model of resident and migrating colon cancer stem cells. iScience. 2024;27(9):110658.

72. Richardson JP, Brown R, Kichik N, Lee S, Priest E, Mogavero S, et al. Candidalysins Are a New Family of Cytolytic Fungal Peptide Toxins. mBio. 2022;13(1):e0351021.

73. Lim SJ, Mohamad Ali MS, Sabri S, Muhd Noor ND, Salleh AB, Oslan SN. Opportunistic yeast pathogen Candida spp.: Secreted and membrane-bound virulence factors. Med Mycol. 2021;59(12):1127–44.

74. Watkins RR, Gowen R, Lionakis MS, Ghannoum M. Update on the Pathogenesis, Virulence, and Treatment of Candida auris. Pathog Immun. 2022;7(2):46–65.

75. Gravdal K, Halvorsen OJ, Haukaas SA, Akslen LA. A switch from E-cadherin to N-cadherin expression indicates epithelial to mesenchymal transition and is of strong and independent importance for the progress of prostate cancer. Clin Cancer Res. 2007;13(23):7003–11.

76. Krisanaprakornkit S, Iamaroon A. Epithelial-mesenchymal transition in oral squamous cell carcinoma. ISRN Oncol. 2012;2012:681469.

77. Nguyen PT, Kudo Y, Yoshida M, Kamata N, Ogawa I, Takata T. N-cadherin expression is involved in malignant behavior of head and neck cancer in relation to epithelial-mesenchymal transition. Histol Histopathol. 2011;26(2):147–56.

78. Pal M, Bhattacharya S, Kalyan G, Hazra S. Cadherin profiling for therapeutic interventions in Epithelial Mesenchymal Transition (EMT) and tumorigenesis. Exp Cell Res. 2018;368(2):137–46.

79. Prudkin L, Liu DD, Ozburn NC, Sun M, Behrens C, Tang X, et al. Epithelial-to-mesenchymal transition in the development and progression of adenocarcinoma and squamous cell carcinoma of the lung. Mod Pathol. 2009;22(5):668–78.

80. Wei J, Wu L, Yang S, Zhang C, Feng L, Wang M, et al. E-cadherin to N-cadherin switching in the TGF-beta1 mediated retinal pigment epithelial to mesenchymal transition. Exp Eye Res. 2022;220:109085.

81. Batlle E, Sancho E, Franci C, Dominguez D, Monfar M, Baulida J, et al. The transcription factor snail is a repressor of E-cadherin gene expression in epithelial tumour cells. Nat Cell Biol. 2000;2(2):84–9.

82. Cano A, Perez-Moreno MA, Rodrigo I, Locascio A, Blanco MJ, del Barrio MG, et al. The transcription factor snail controls epithelial-mesenchymal transitions by repressing E-cadherin expression. Nat Cell Biol. 2000;2(2):76–83.

83. Ikenouchi J, Matsuda M, Furuse M, Tsukita S. Regulation of tight junctions during the epithelium-mesenchyme transition: direct repression of the gene expression of claudins/occludin by Snail. J Cell Sci. 2003;116(Pt 10):1959–67.

84. Martinez-Estrada OM, Culleres A, Soriano FX, Peinado H, Bolos V, Martinez FO, et al. The transcription factors Slug and Snail act as repressors of Claudin-1 expression in epithelial cells. Biochem J. 2006;394(Pt 2):449–57.

85. Porta-de-la-Riva M, Stanisavljevic J, Curto J, Franci C, Diaz VM, Garcia de Herreros A, et al. TFCP2c/LSF/LBP-1c is required for Snail1-induced fibronectin gene expression. Biochem J. 2011;435(3):563–8.

86. Stanisavljevic J, Porta-de-la-Riva M, Batlle R, de Herreros AG, Baulida J. The p65 subunit of NF-kappaB and PARP1 assist Snail1 in activating fibronectin transcription. J Cell Sci. 2011;124(Pt 24):4161–71.

87. Nieto MA, Huang RY, Jackson RA, Thiery JP. Emt: 2016. Cell. 2016;166(1):21–45.

88. Guo HB, Johnson H, Randolph M, Pierce M. Regulation of homotypic cell-cell adhesion by branched N-glycosylation of N-cadherin extracellular EC2 and EC3 domains. J Biol Chem. 2009;284(50):34986–97.

89. Cummings RD, Kornfeld S. Characterization of the structural determinants required for the high affinity interaction of asparagine-linked oligosaccharides with immobilized Phaseolus vulgaris leukoagglutinating and erythroagglutinating lectins. J Biol Chem. 1982;257(19):11230–4.

90. Tulsiani DR, Harris TM, Touster O. Swainsonine inhibits the biosynthesis of complex glycoproteins by inhibition of Golgi mannosidase II. J Biol Chem. 1982;257(14):7936–9.

91. de-Freitas-Junior JC, Bastos LG, Freire-Neto CA, Rocher BD, Abdelhay ES, Morgado-Diaz JA. N-glycan biosynthesis inhibitors induce in vitro anticancer activity in colorectal cancer cells. J Cell Biochem. 2012;113(9):2957–66.

92. Sima LE, Matei D, Condello S. The Outside-In Journey of Tissue Transglutaminase in Cancer. Cells. 2022;11(11).

93. Narunsky-Haziza L, Sepich-Poore GD, Livyatan I, Asraf O, Martino C, Nejman D, et al. Pan-cancer analyses reveal cancer-type-specific fungal ecologies and bacteriome interactions. Cell. 2022;185(20):3789–806 e17.

94. Cellura D, Pickard K, Quaratino S, Parker H, Strefford JC, Thomas GJ, et al. miR-19-Mediated Inhibition of Transglutaminase-2 Leads to Enhanced Invasion and Metastasis in Colorectal Cancer. Mol Cancer Res. 2015;13(7):1095–105.

95. Delaine-Smith R, Wright N, Hanley C, Hanwell R, Bhome R, Bullock M, et al. Transglutaminase-2 Mediates the Biomechanical Properties of the Colorectal Cancer Tissue Microenvironment that Contribute to Disease Progression. Cancers (Basel). 2019;11(5).

96. Allert S, Forster TM, Svensson CM, Richardson JP, Pawlik T, Hebecker B, et al. Candida albicans-Induced Epithelial Damage Mediates Translocation through Intestinal Barriers. mBio. 2018;9(3).

97. Koh AY, Kohler JR, Coggshall KT, Van Rooijen N, Pier GB. Mucosal damage and neutropenia are required for Candida albicans dissemination. PLoS Pathog. 2008;4(2):e35.

98. Canaparo R, Foglietta F, Pepa CD, Serpe L. Spotlight on membrane fluidity of normal and cancer cells: Implications for cancer diagnosis and treatment. Eur J Pharmacol. 2025;1006:178152.

99. Shimolina L, Efremov YM, Gulin A, Ignatova N, Aybush A, Kuimova MK, et al. Comparison of viscoelastic properties of breast cancer and normal cells using AFM, FLIM and ToF-SIMS techniques. Sci Rep. 2025;16(1):1655.

100. Lachat J, Pascault A, Thibaut D, Le Borgne R, Verbavatz JM, Weiner A. Trans-cellular tunnels induced by the fungal pathogen Candida albicans facilitate invasion through successive epithelial cells without host damage. Nat Commun. 2022;13(1):3781.

101. Hernandez R, Rupp S. Human epithelial model systems for the study of Candida infections in vitro: part II. Histologic methods for studying fungal invasion. Methods Mol Biol. 2009;470:105–23.

102. Sohn K, Senyurek I, Fertey J, Konigsdorfer A, Joffroy C, Hauser N, et al. An in vitro assay to study the transcriptional response during adherence of Candida albicans to different human epithelia. FEMS Yeast Res. 2006;6(7):1085–93.

103. Weide MR, Ernst JF. Caco-2 monolayer as a model for transepithelial migration of the fungal pathogen Candida albicans. Mycoses. 1999;42 Suppl 2:61–7.

104. Kalinin A, Marekov LN, Steinert PM. Assembly of the epidermal cornified cell envelope. J Cell Sci. 2001;114(Pt 17):3069–70.

105. Nemes Z, Steinert PM. Bricks and mortar of the epidermal barrier. Exp Mol Med. 1999;31(1):5–19.

106. Silva S, Hooper SJ, Henriques M, Oliveira R, Azeredo J, Williams DW. The role of secreted aspartyl proteinases in Candida tropicalis invasion and damage of oral mucosa. Clin Microbiol Infect. 2011;17(2):264–72.

107. Yu S, Li W, Liu X, Che J, Wu Y, Lu J. Distinct Expression Levels of ALS, LIP, and SAP Genes in Candida tropicalis with Diverse Virulent Activities. Front Microbiol. 2016;7:1175.

108. Lu S, Saydak M, Gentile V, Stein JP, Davies PJ. Isolation and characterization of the human tissue transglutaminase gene promoter. J Biol Chem. 1995;270(17):9748–56.

109. Prall F. Tumour budding in colorectal carcinoma. Histopathology. 2007;50(1):151–62.

110. Zlobec I, Lugli A. Invasive front of colorectal cancer: dynamic interface of pro-/anti-tumor factors. World J Gastroenterol. 2009;15(47):5898–906.

111. Haerinck J, Goossens S, Berx G. The epithelial-mesenchymal plasticity landscape: principles of design and mechanisms of regulation. Nat Rev Genet. 2023.

112. Mullins RDZ, Pal A, Barrett TF, Heft Neal ME, Puram SV. Epithelial-Mesenchymal Plasticity in Tumor Immune Evasion. Cancer Res. 2022;82(13):2329–43.

113. Frank CF, Hostetter MK. Cleavage of E-cadherin: a mechanism for disruption of the intestinal epithelial barrier by Candida albicans. Transl Res. 2007;149(4):211–22.

114. Villar CC, Kashleva H, Nobile CJ, Mitchell AP, Dongari-Bagtzoglou A. Mucosal tissue invasion by Candida albicans is associated with E-cadherin degradation, mediated by transcription factor Rim101p and protease Sap5p. Infect Immun. 2007;75(5):2126–35.

115. Leone L, Mazzetta F, Martinelli D, Valente S, Alimandi M, Raffa S, et al. Klebsiella pneumoniae Is Able to Trigger Epithelial-Mesenchymal Transition Process in Cultured Airway Epithelial Cells. PLoS One. 2016;11(1):e0146365.

116. Liu W, Lu JY, Wang YJ, Xu XX, Chen YC, Yu SX, et al. Vaccinia virus induces EMT-like transformation and RhoA-mediated mesenchymal migration. J Med Virol. 2023;95(8):e29041.

117. Sun M, Peng Z, Shen W, Guo X, Liao Y, Huang Y, et al. Synergism of Fusobacterium periodonticum and N-nitrosamines promote the formation of EMT subtypes in ESCC by modulating Wnt3a palmitoylation. Gut Microbes. 2024;16(1):2391521.

118. Zhang R, Takigawa H, Maruyama H, Nambu T, Mashimo C, Okinaga T. Fisetin Inhibits Periodontal Pathogen-Induced EMT in Oral Squamous Cell Carcinoma via the Wnt/beta-Catenin Pathway. Nutrients. 2025;17(22).

119. Jiang H, Xu B. The critical role of epithelial-mesenchymal transition (EMT) in colorectal cancer progression and therapeutic outcomes. Crit Rev Oncol Hematol. 2026;220:105189.

120. Onder TT, Gupta PB, Mani SA, Yang J, Lander ES, Weinberg RA. Loss of E-cadherin promotes metastasis via multiple downstream transcriptional pathways. Cancer Res. 2008;68(10):3645–54.

121. Zhao Y, Sato Y, Isaji T, Fukuda T, Matsumoto A, Miyoshi E, et al. Branched N-glycans regulate the biological functions of integrins and cadherins. FEBS J. 2008;275(9):1939–48.

122. de-Souza-Ferreira M, Ferreira EE, de-Freitas-Junior JCM. Aberrant N-glycosylation in cancer: MGAT5 and beta1,6-GlcNAc branched N-glycans as critical regulators of tumor development and progression. Cell Oncol (Dordr). 2023;46(3):481–501.

123. Takahashi M, Hasegawa Y, Maeda K, Kitano M, Taniguchi N. Role of glycosyltransferases in carcinogenesis; growth factor signaling and EMT/MET programs. Glycoconj J. 2022;39(2):167–76.

124. Vajaria BN, Patel PS. Glycosylation: a hallmark of cancer? Glycoconj J. 2017;34(2):147–56.

125. Holm M, Nummela P, Heiskanen A, Satomaa T, Kaprio T, Mustonen H, et al. N-glycomic profiling of colorectal cancer according to tumor stage and location. PLoS One. 2020;15(6):e0234989.

126. Venkitachalam S, Revoredo L, Varadan V, Fecteau RE, Ravi L, Lutterbaugh J, et al. Biochemical and functional characterization of glycosylation-associated mutational landscapes in colon cancer. Sci Rep. 2016;6:23642.

127. Huang HC, Huang YL, Chen YJ, Wu HY, Hsu CL, Kao HF, et al. The branched N-glycan of PD-L1 predicts immunotherapy responses in patients with recurrent/metastatic HNSCC. Oncogenesis. 2024;13(1):36.

128. Nie H, Saini P, Miyamoto T, Liao L, Zielinski RJ, Liu H, et al. Targeting branched N-glycans and fucosylation sensitizes ovarian tumors to immune checkpoint blockade. Nat Commun. 2024;15(1):2853.

129. Silva MC, Fernandes A, Oliveira M, Resende C, Correia A, de-Freitas-Junior JC, et al. Glycans as Immune Checkpoints: Removal of Branched N-glycans Enhances Immune Recognition Preventing Cancer Progression. Cancer Immunol Res. 2020;8(11):1407–25.

130. Gornik O, Royle L, Harvey DJ, Radcliffe CM, Saldova R, Dwek RA, et al. Changes of serum glycans during sepsis and acute pancreatitis. Glycobiology. 2007;17(12):1321–32.

131. Heindel DW, Chen S, Aziz PV, Chung JY, Marth JD, Mahal LK. Glycomic Analysis Reveals a Conserved Response to Bacterial Sepsis Induced by Different Bacterial Pathogens. ACS Infect Dis. 2022;8(5):1075–85.

132. Lu LL, Das J, Grace PS, Fortune SM, Restrepo BI, Alter G. Antibody Fc Glycosylation Discriminates Between Latent and Active Tuberculosis. J Infect Dis. 2020;222(12):2093–102.

133. Muenchhoff M, Chung AW, Roider J, Dugast AS, Richardson S, Kloverpris H, et al. Distinct Immunoglobulin Fc Glycosylation Patterns Are Associated with Disease Nonprogression and Broadly Neutralizing Antibody Responses in Children with HIV Infection. mSphere. 2020;5(6).

134. Qin R, Mahal LK. The host glycomic response to pathogens. Curr Opin Struct Biol. 2021;68:149–56.

135. Torpy H, Chau TH, Chatterjee S, Chernykh A, Torpy DJ, Meyer EJ, et al. Impact of different pathogen classes on the serum N-glycome in septic shock. BBA Adv. 2025;7:100138.

136. Lee KN, Maxwell MD, Patterson MK, Jr., Birckbichler PJ, Conway E. Identification of transglutaminase substrates in HT29 colon cancer cells: use of 5-(biotinamido)pentylamine as a transglutaminase-specific probe. Biochim Biophys Acta. 1992;1136(1):12–6.

137. Kang S, Oh SC, Min BW, Lee DH. Transglutaminase 2 Regulates Self-renewal and Stem Cell Marker of Human Colorectal Cancer Stem Cells. Anticancer Res. 2018;38(2):787–94.

138. Condello S, Prasad M, Atwani R, Matei D. Tissue transglutaminase activates integrin-linked kinase and beta-catenin in ovarian cancer. J Biol Chem. 2022;298(8):102242.

139. Kumar A, Xu J, Brady S, Gao H, Yu D, Reuben J, et al. Tissue transglutaminase promotes drug resistance and invasion by inducing mesenchymal transition in mammary epithelial cells. PLoS One. 2010;5(10):e13390.

140. Mehta K, Kumar A, Kim HI. Transglutaminase 2: a multi-tasking protein in the complex circuitry of inflammation and cancer. Biochem Pharmacol. 2010;80(12):1921–9.

141. Tempest R, Guarnerio S, Maani R, Cooper J, Peake N. The Biological and Biomechanical Role of Transglutaminase-2 in the Tumour Microenvironment. Cancers (Basel). 2021;13(11).

142. Griffiths JS, Camilli G, Kotowicz NK, Ho J, Richardson JP, Naglik JR. Role for IL-1 Family Cytokines in Fungal Infections. Front Microbiol. 2021;12:633047.

143. Naglik JR, Konig A, Hube B, Gaffen SL. Candida albicans-epithelial interactions and induction of mucosal innate immunity. Curr Opin Microbiol. 2017;40:104–12.

144. Baram T, Rubinstein-Achiasaf L, Ben-Yaakov H, Ben-Baruch A. Inflammation-Driven Breast Tumor Cell Plasticity: Stemness/EMT, Therapy Resistance and Dormancy. Front Oncol. 2020;10:614468.

145. Krug J, Rodrian G, Petter K, Yang H, Khoziainova S, Guo W, et al. N-glycosylation Regulates Intrinsic IFN-gamma Resistance in Colorectal Cancer: Implications for Immunotherapy. Gastroenterology. 2023;164(3):392–406 e5.

