## Supplementary figures and images for "*Candida albicans* promotes self-adhesion and tumor cell progression of colorectal cancer cells"

### Supplemental figures S1-S8

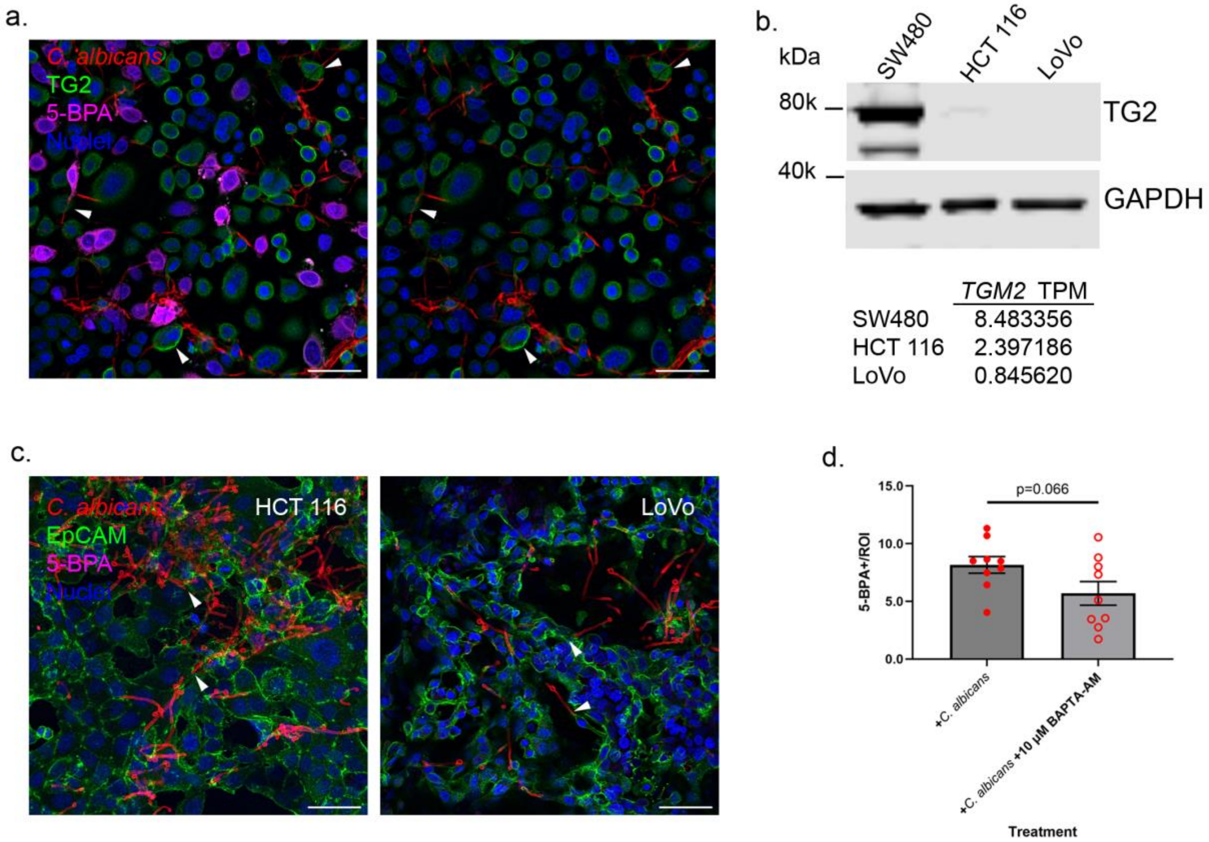

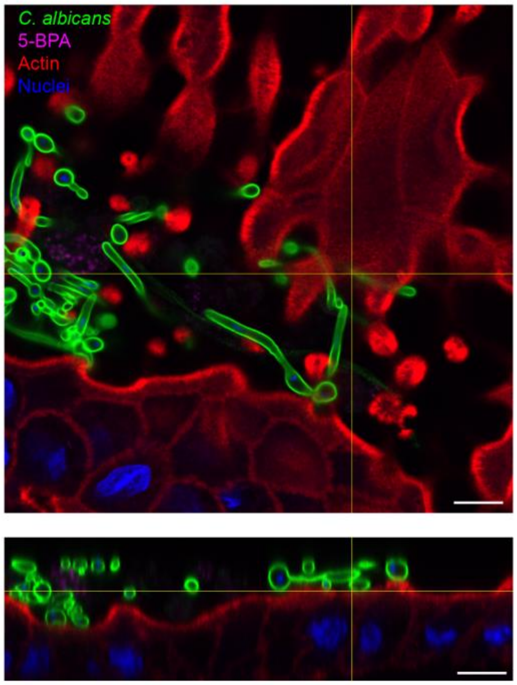

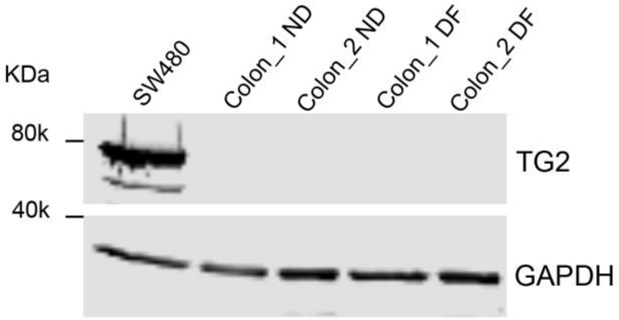

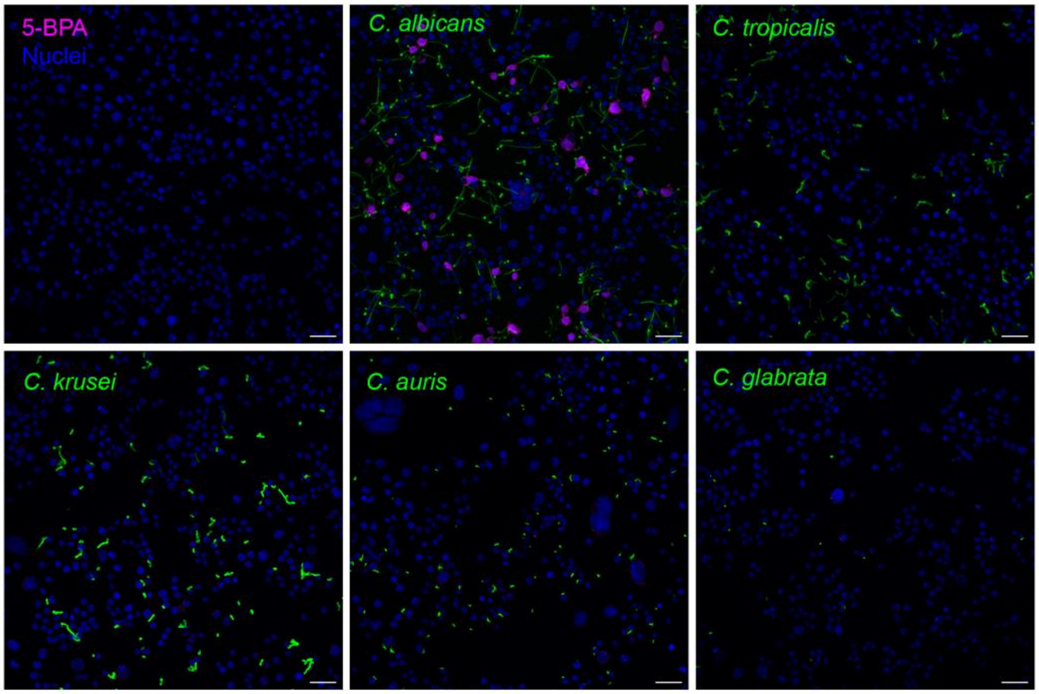

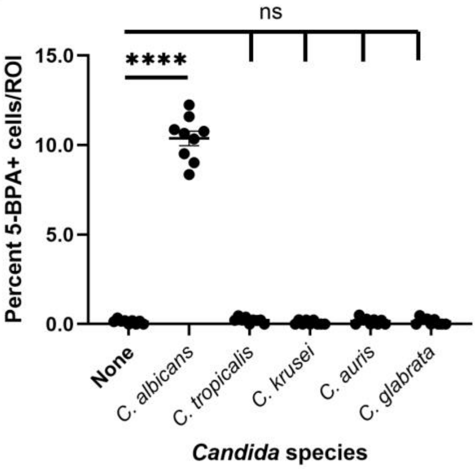

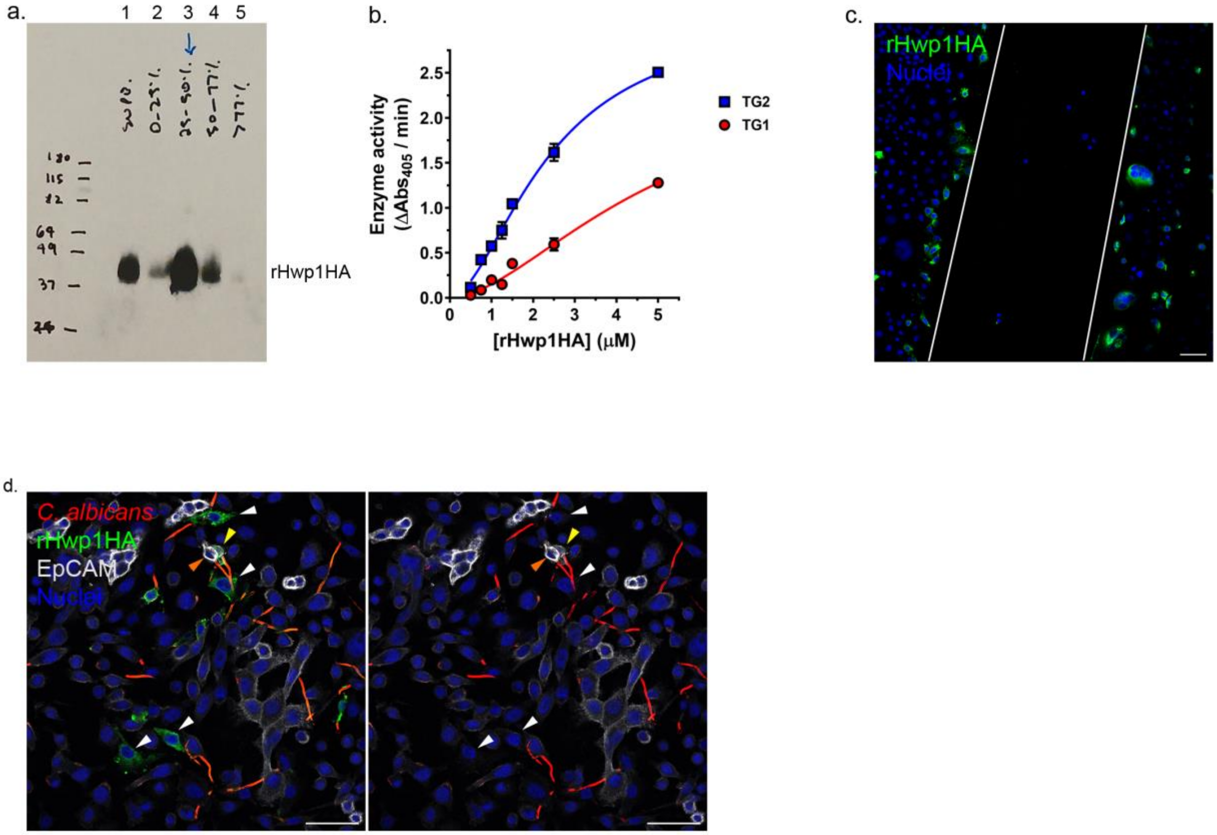

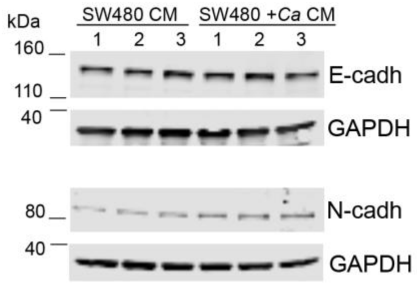

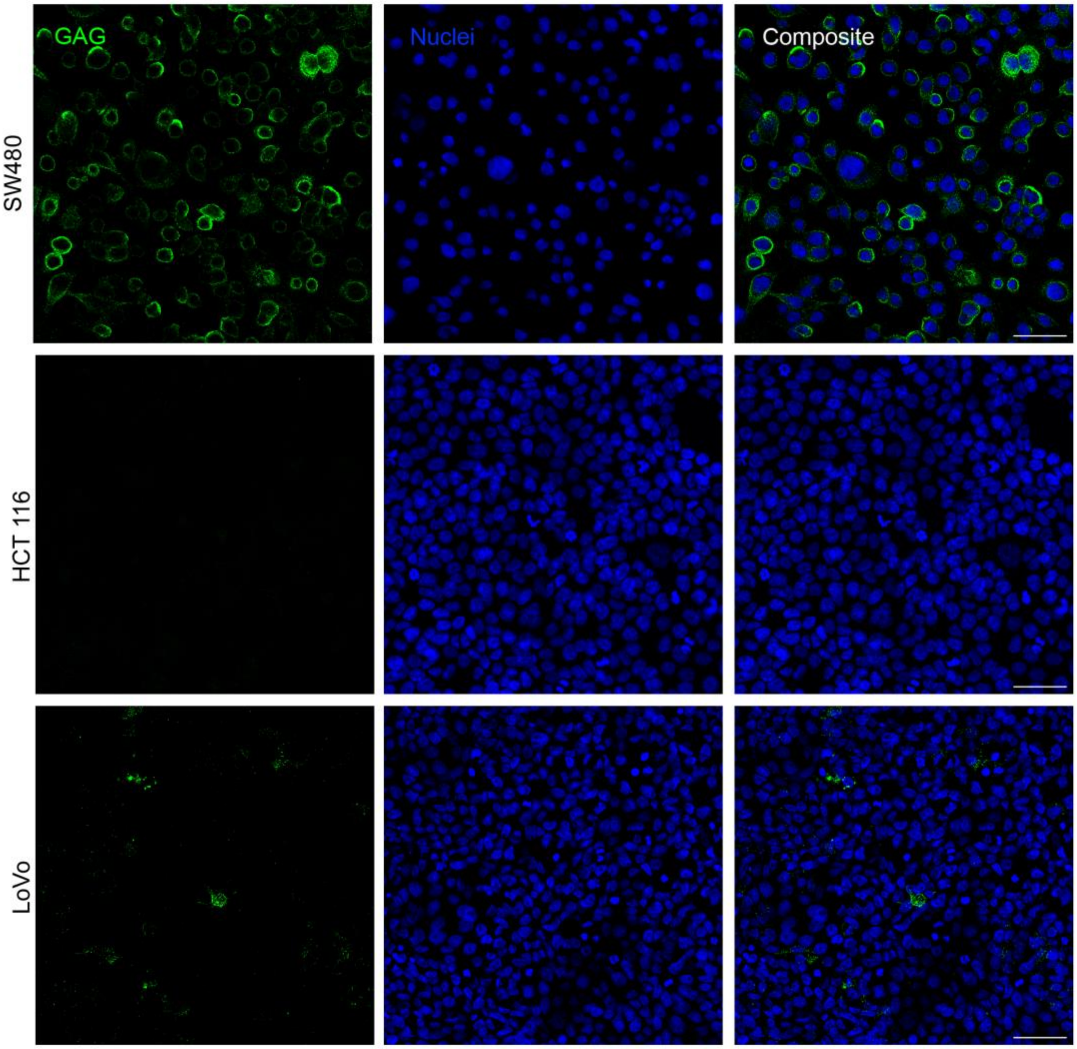

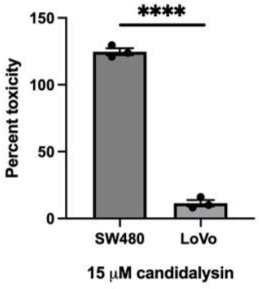

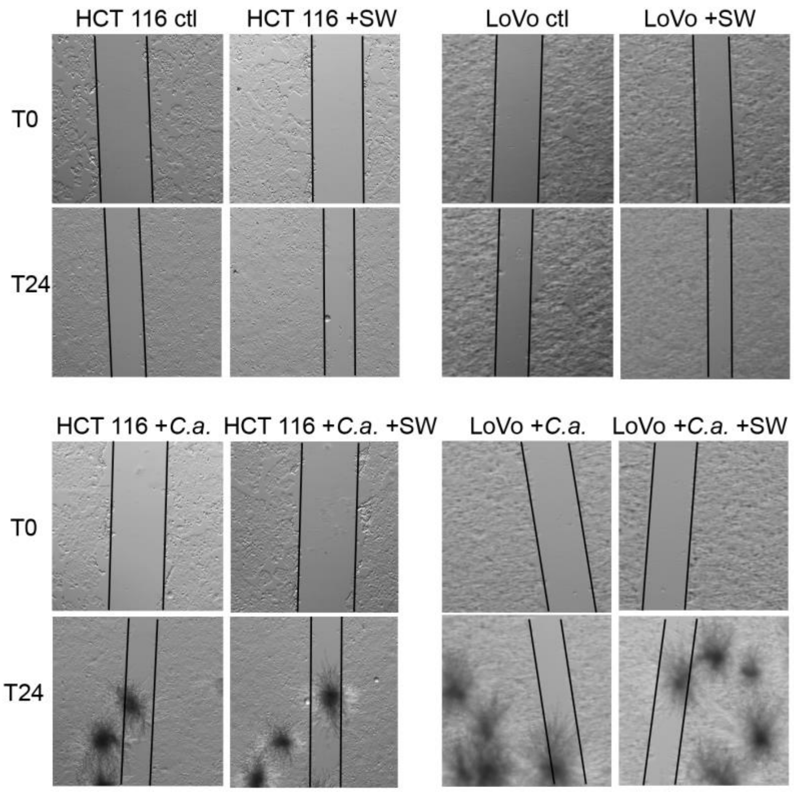

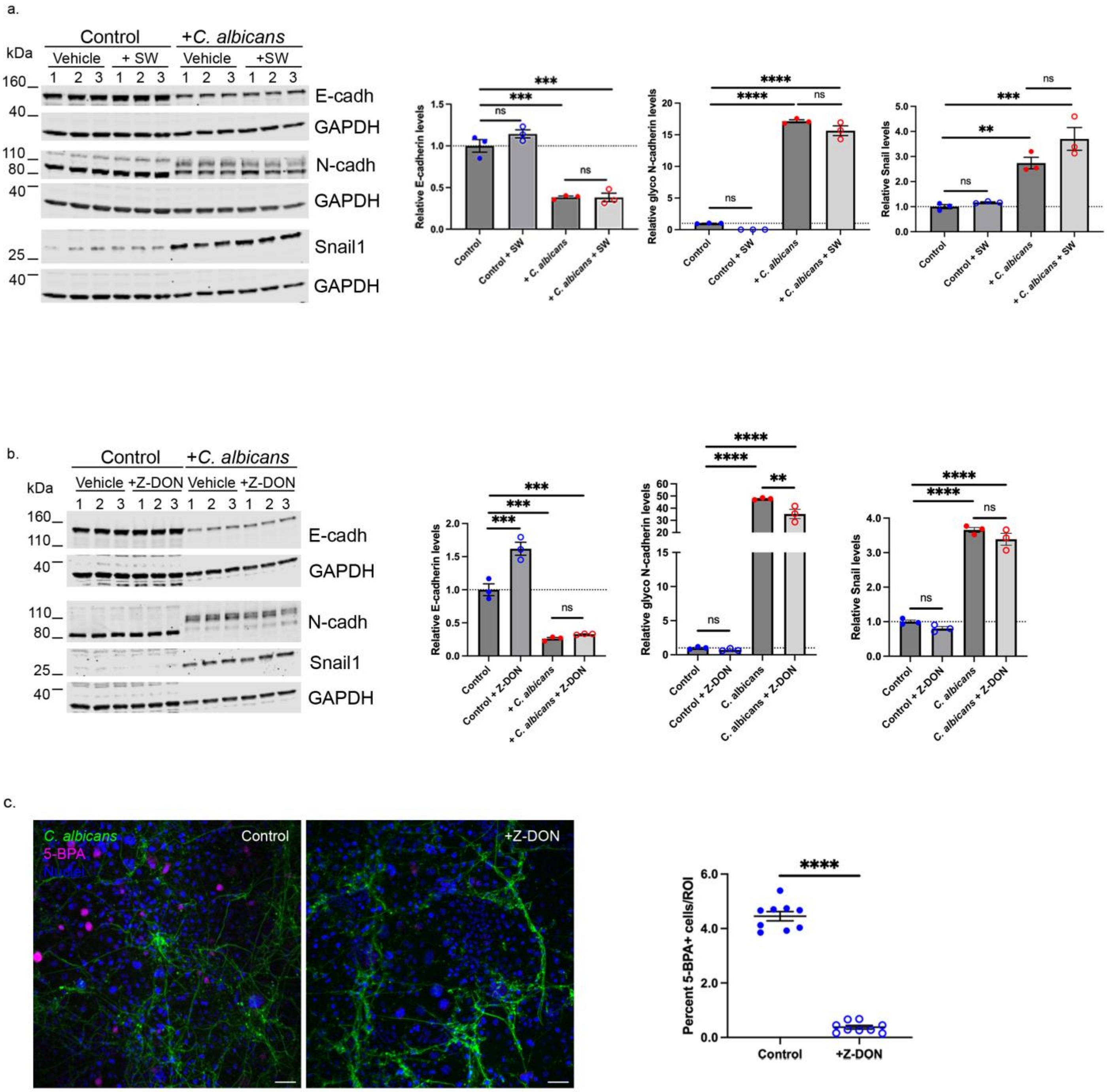

Fig. 4a

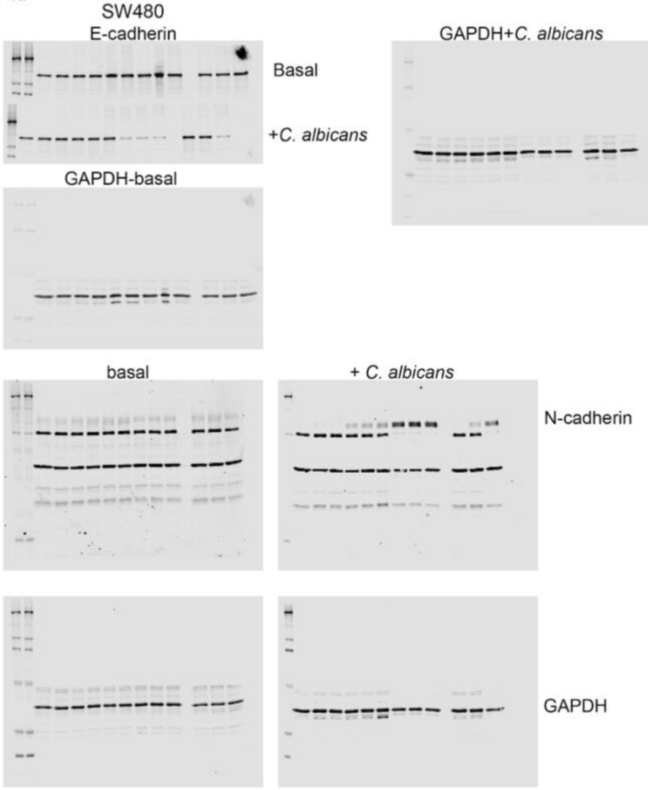


Fig. 4c

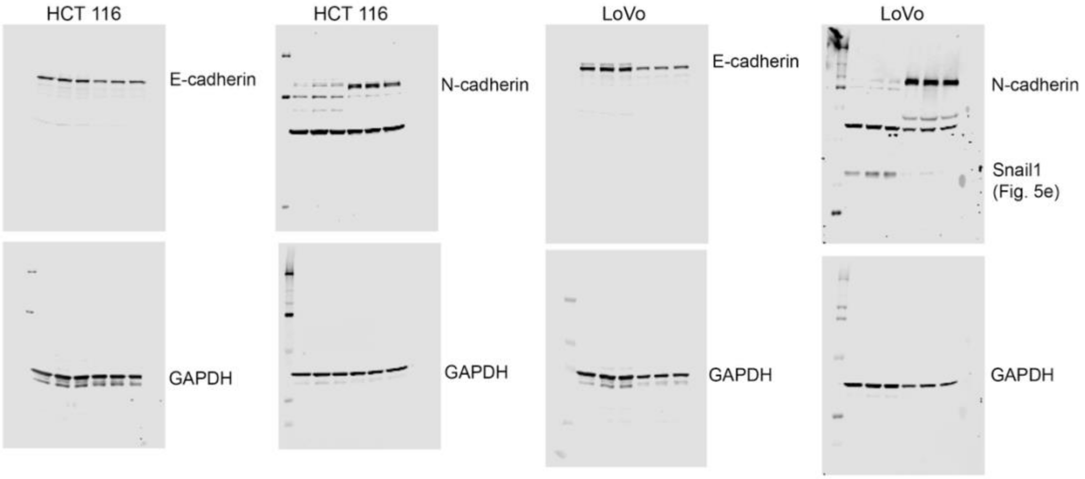


Fig. 4f

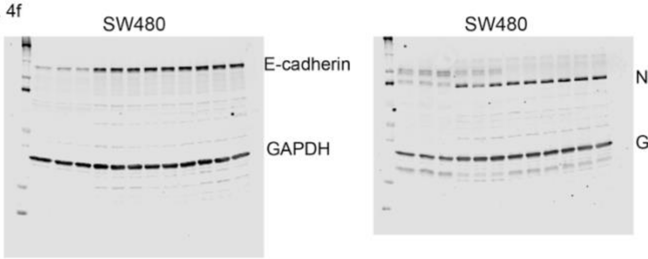

Fig. 5a

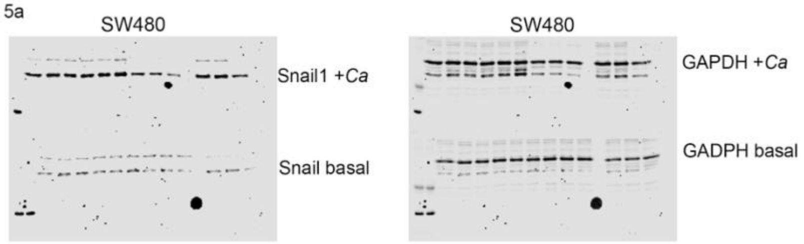


Fig. 5c

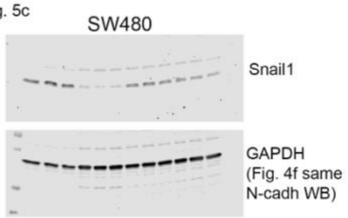


Fig. 5e

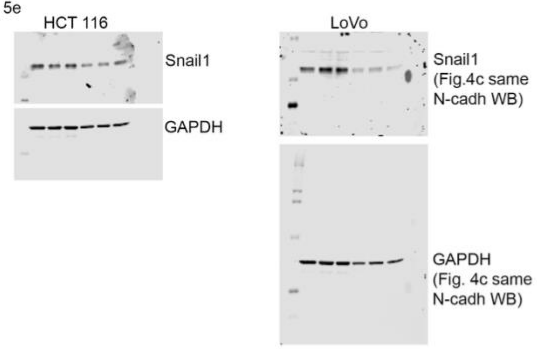


Fig. 6a

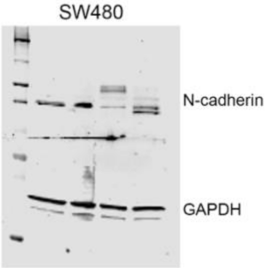


Fig. 6b

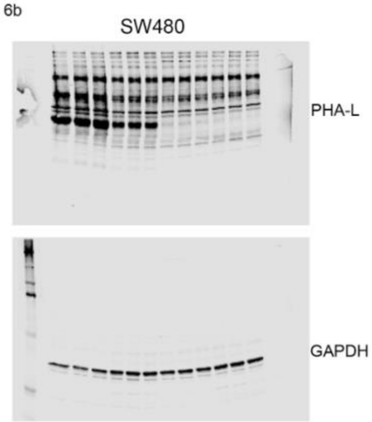


Fig. 6d

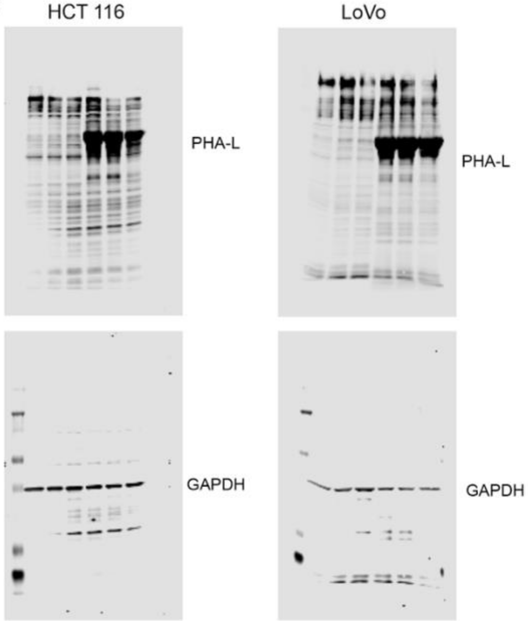

Fig. S1

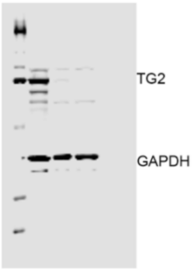


Fig. S2

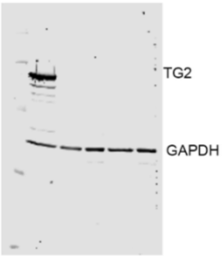


Fig. S5a

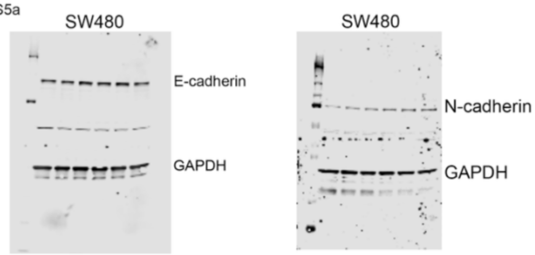


Fig. S7a

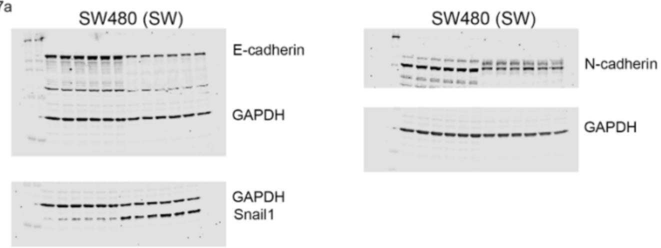


Fig. S7b

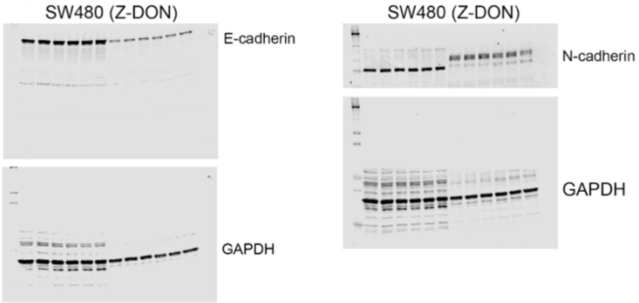
